# Human POT1 Unfolds G-Quadruplexes by Conformational Selection

**DOI:** 10.1101/691048

**Authors:** Jonathan B. Chaires, Robert D. Gray, William L. Dean, Robert Monsen, Lynn W. DeLeeuw, Vilius Stribinskis, John O. Trent

**Affiliations:** James Graham Brown Cancer Center, University of Louisville, 505 S. Hancock St., Louisville, KY 40202 USA

## Abstract

The reaction mechanism by which shelterin protein POT1 (<u>P</u>rotection <u>o</u>f <u>T</u>elomeres) unfolds human telomeric G-quadruplex structures is not fully understood. We report here kinetic, thermodynamic, hydrodynamic and computational studies that show that a conformational selection mechanism, in which POT1 binding is coupled to an obligatory unfolding reaction, is the most plausible mechanism. We show that binding of the single-strand oligonucleotide d[TTAGGGTTAG] to POT1 is fast, with an apparent relaxation time of 80.0 ± 0.4 ms, and strong, with a binding free energy of −10.1 ± 0.3 kcal mol^−1^. That favourable free energy arises from a large favourable enthalpy contribution of −38.2 ± 0.3 kcal mol^−1^. In sharp contrast, the binding of POT1 to an initially folded 24 nt G-quadruplex structure is slow, with an average relaxation time of 2000-3000 s. Fluorescence, circular dichroism and analytical ultracentrifugation studies show that POT1 binding is coupled to quadruplex unfolding with a final stoichiometry of 2 POT1 molecules bound per 24 nt DNA. The binding isotherm for the POT1-quadruplex binding interaction is sigmoidal, indicative of a complex reaction. A conformational selection model that includes equilibrium constants for both G-quadruplex unfolding and POT1 binding to the resultant single-strand provides an excellent quantitative fit to the experimental binding data. The overall favourable free energy of the POT1-quadruplex interaction is −7.1 kcal mol^−1^, which arises from a balance between unfavourable free energy of +3.4 kcal mol^−1^ for quadruplex unfolding and a large, favorable free energy of −10.5 kcal mol^−1^ for POT1 binding. We show that POT1 can unfold and bind to any conformational form of human telomeric G-quadruplex (antiparallel, hybrid or parallel), but will not interact with duplex DNA or with a parallel G-quadruplex structure formed by a c-myc promoter sequence. Finally, molecular dynamics simulations provide a detailed structural model of a 2:1 POT1:DNA complex that is fully consistent with experimental biophysical results.

## INTRODUCTION

POT1 (<u>P</u>rotection <u>O</u>f <u>T</u>elomeres 1) is a telomere single-stranded DNA binding protein found in a variety of organisms(1–4) that is essential for telomerase activity (5,6). POT1 is an integral component of the shelterin complex (7,8) and is the only protein in the complex to bind with high sequence specificity to the G-rich ssDNA 3’-overhang. It does not bind to double stranded telomeric DNA or to the complementary C-rich strand (4). Human POT1 has two functional domains: an N-terminal oligonucleotide-binding (OB) domain that is required for DNA binding and a C-terminal domain that binds the shelterin protein TPP1. In humans, POT1 is involved in regulation of telomere length and telomerase activity (5,6). *In vitro*, POT1 inhibits telomerase activity at the 3’-end of DNA by controlling accessibility of the single-stranded DNA substrate to telomerase (9). Although the exact role of the DNA binding activity of POT1 is unknown, the crystal structure of human POT1 with the optimal telomeric DNA sequence implies that it physically caps the end of chromosomes by sequestering the last guanine base of DNA into a hydrophobic pocket, making it inaccessible to telomerase (4,10). POT1 capping of the chromosome ends may also prevent the ends from eliciting a DNA damage response (10). However, *in vitro* binding of shelterin protein TPP1 to POT1 relieves telomerase inhibition and increases the repeat addition processivity of the enzyme (9). POT1 also acts as a positive regulator of telomere length as overexpression of full-length POT1 in telomerase positive cells leads to the lengthening of telomeres (11). POT1 has also been implicated in binding within the hypothetical D-loop part of the proposed T-loop configuration of telomeres (12).

### POT1-ssDNA Structure

A high resolution crystal structure of human POT1 bound to the optimal human telomeric binding sequence d[TAGGGTTAG] was reported by the Cech group in 2004 (10). This structure reveals two N-terminal OB domains spanning the length of the bound oligonucleotide. It also can be inferred from the crystal structure that with minimal movement of the terminal bases, longer sequences could be bound by multiple POT1 N-terminal OB domains. There are no high-resolution structures of POT1 bound to long single-stranded telomeric sequences. However, Taylor and coworkers (13) examined binding of POT1 to 72-144 nt tracts of DNA by electrophoretic mobility shift assays, size-exclusion chromatography and electron microscopy (EM). They found that the ssDNA was fully occupied and coated by POT1 and its variants, with one POT1 bound consecutively to every 12 nt repeat. EM showed that a 144 nt DNA saturated with 12 POT1-TPP1 heterodimers formed ordered, compact assemblies. Similar structures were seen for POT1 alone bound to a 132 nt ssDNA.

### Mechanism of POT1 unfolding of telomeric quadruplexes

The mechanism by which POT1 binds to, and unfolds, quadruplex DNA is not fully understood. Early studies of POT1 DNA interactions were limited to short single-stranded oligonucleotides for which binding was uncoupled from any unfolding process (5,14). Several subsequent studies have addressed the unfolding of G4 DNA by POT1. First, the Cech laboratory showed that human POT1 disrupts G4s and speculated that “hPOT1 may function simply by trapping the unfolded forms of … telomeric primers in an equilibrium population” (15). They showed that when facilitating telomerase activity, hPOT1 does not act catalytically but formed a stoichiometric complex with the DNA, freeing its 3’ tail. Interestingly, they found that a short antisense oligonucleotide was found to duplicate the effect of POT1 on G4 unfolding and facilitation of telomerase activity.

A second study from the Wang and Opresko laboratories used single-molecule atomic force microscopy (AFM) to monitor POT1 binding to longer telomeric G4 structures and proposed a different model (16). In contrast to our published observations that long telomeric DNA always form structures with the maximal number of quadruplex units (17), they reported that none of the sequences they studied by AFM were fully folded to form the maximal number of G4 units (16). POT1 appeared to bind to single-stranded gaps between quadruplex units. The authors proposed a “steric driver” mechanism in which such binding destabilized adjacent quadruplexes (by an unstated mechanism) to cause unfolding, and argued against the simple static trapping mechanism for POT1 unfolding of G4 structures. POT1 molecules were proposed to slide along the exposed single-strand to stabilize the fully unfolded state. Curiously, the antisense oligonucleotide 5’CCTAACCCTAACC was reported to be less effective than POT1 at disrupting G4 structures, in contrast to the results from the Cech laboratory.

In a third study (by the Opresko and Myong laboratories), single-molecule FRET results indicated that two POT1 molecules bound sequentially, beginning at the 3’ end, to an initially folded telomeric G4 structure (18).

A fourth study by Ray et. al. also used single-molecule FRET (19). These authors reported that POT1 unfolded both the antiparallel telomeric G4 form in Na+ and the “hybrid” form in K+, albeit with different binding stoichiometries of 1 and 2 POT1 molecules per G4, respectively. They reported that POT1 was unable to unfold the parallel telomeric G4 form. They used the phenomenological Hill equation to analyse their binding isotherms, and noted that the estimated binding constant K_eq_ was a composite term that “represents both POT1-mediated GQ unfolding and POT1 binding to the unfolded DNA”.

Finally, a study from the Taylor laboratory (20) used an electrophoretic mobility shift assay to monitor the interaction of a POT1-TPP1 complex with a variety of telomeric G4 structures. A sequential two-step binding model was used to analyse their binding isotherms, and circular dichroism was used to show that the initially folded G4 structure was disrupted upon binding. The authors discussed the complexity of POT1 binding to an initially folded G4 structure, suggesting the possibility that “… POT1 binds first to one of the TTA loops, and this interaction weakens the G-quadruplex hydrogen bonding network.” allowing an addition POT1 to bind. Or, they noted, “[alternatively, POT1 - TPP1 binding may simply trap the unfolded DNA conformation. In this model, the unfolding of the G-quadruplex occurs in a pre-equilibrium step that is followed by POT1 - TPP1 binding that prevents refolding”. They were not able to decide definitively which of these mechanisms was most likely.

Collectively these studies show that POT1 unfolds G4 DNA to produce a single-strand DNA-POT1 complex, but that there is no consensus for the mechanism by which it does so. The thermodynamic linkage between G4 unfolding and POT1 binding has not yet been quantitatively explained.

### Goals of this study

The primary goal of our study is to understand the mechanism by which POT1 unfolds telomeric G4 structures. In particular, we wish to understand the coupling and thermodynamic linkage between G4 unfolding and POT1 binding. Our study builds on our previous experimental studies of the kinetics and thermodynamics of the folding and unfolding of telomeric G4 structures (21–30). We use a battery of biophysical tools to probe the kinetics and thermodynamics of POT1 binding to both the short single-stranded telomeric DNA recognition sequence and to longer, initially folded, telomeric G4 sequences. Our results show that a conformational-selection binding model in which POT1 binding is coupled to an obligatory unfolding step of the G4 structure is most consistent with our data. In addition, we used molecular dynamics simulations to build a detailed model for the POT1-DNA complex that predicts experimentally accessible biophysical properties that are fully consistent with our biophysical results.

## Results and Analysis

### Binding of POT1 to a single-stranded DNA oligonucleotide is strong and fast

We studied the equilibrium binding of the single-stranded oligonucleotide d[TTAGGGTTAG] (O1) to POT1 by several spectroscopic methods and by ITC. Figure 1A shows binding isotherms obtained by using either FRET labeled O1 or intrinsic POT1 tryptophan fluorescence. These data were fit to a 1:1 binding model that yielded estimates for the dissociation constant K_d_ of 26.4 and 29.6 nM, respectively. Additional titration experiments, done using microscale thermophoresis or by fluorescence with O1 labeled with 2-aminopurine (Supplementary Figure S1), yielded similar K_d_ estimates as summarized in Supplementary Table S2. Figure 1B shows the results of an ITC binding experiment. A 1:1 binding model fits the data, yielding K_d_ = 59.4 nM, ΔH_b_ = −38.2 kcal mol^−1^ and n=0.67. The collective binding data (Table S2, Figure 1) are in good agreement. The thermodynamic profile determined for the POT1-O1 binding interaction from these data is ΔG = −10.1 ±0.3 kcal mol^−1^, ΔH_b_ = −38.2±0.3 kcal mol^−1^ and −TΔS = +23.2±0.4 kcal mol^−1^, as shown in the inset in Figure 1B. The favorable free energy of binding arises from the large favorable enthalpy contribution, and is opposed by entropy.

**Figure 1.**
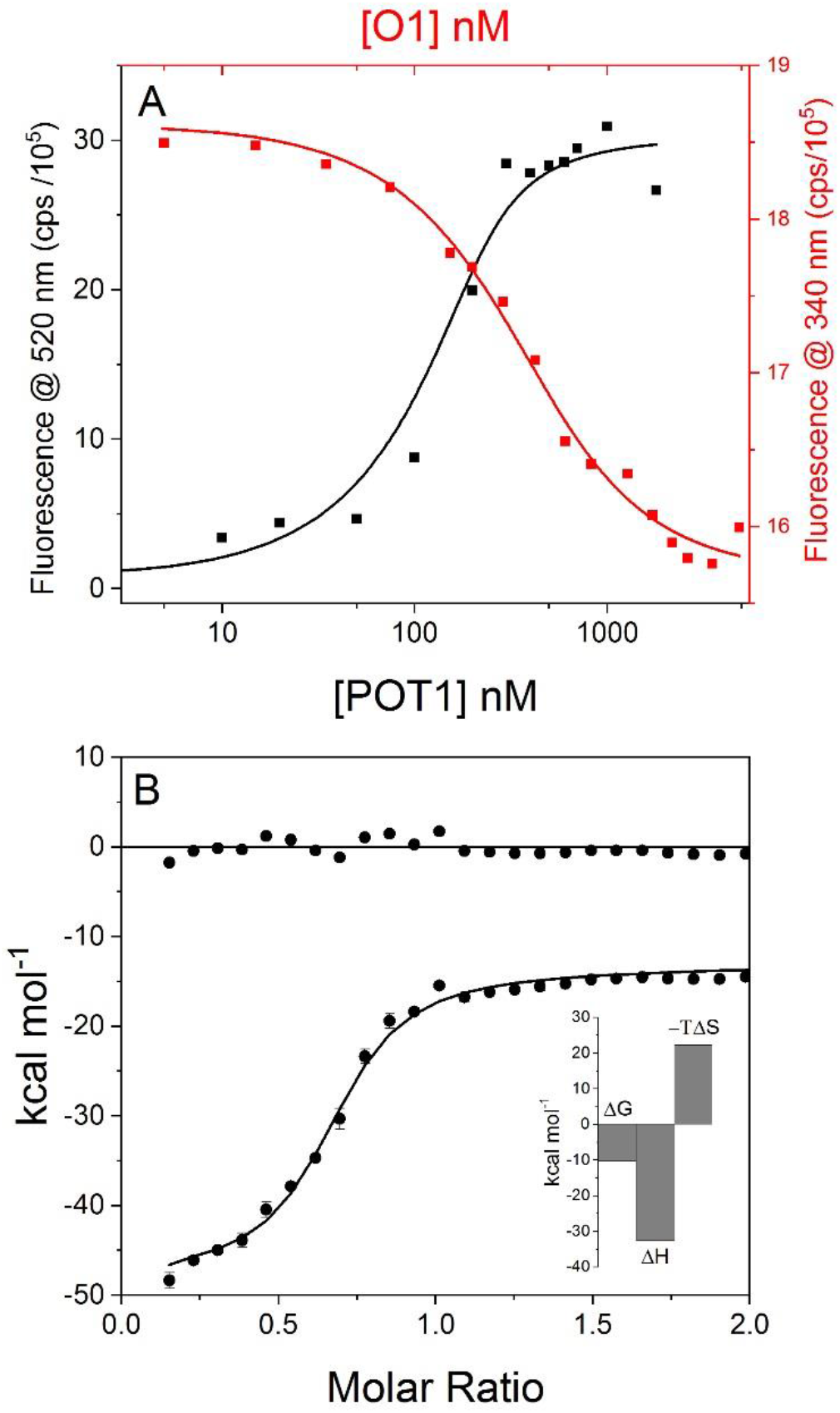
Binding of the deoxyoligonucleotide d[TTAGGGTTAG] (“O1”) to POT1. **(A)** Isothermal titration experiments monitored by changes in FRET-labeled O1 fluorescence (black) or changes in intrinsic POT1 tryptophan fluorescence (red). In the first case, a fixed concentration of O1 was titrated with increasing amounts of POT1. In the second case, a fixed concentration of POT1 was titrated with increasing amounts of unlabeled O1. The lines are the best fits to a 1:1 binding model obtained by nonlinear least-squares fitting of the data, with the results shown in Table S2 and discussed in the text. (**B**). Isothermal titration calorimetry results for the titration of O1 into a POT1 solution. The solid lines show the best fit to a 1:1 binding model, with the residuals (data – fit) shown at the top of the panel centered around 0. The fitted parameters are reported in the text and in Table S2. The inset shows the thermodynamic profile for O1 binding to POT1, showing that the favorable binding free energy (ΔG = −10.1 kcal mol^−1^) results from the difference between a favorable binding enthalpy (ΔH = −33.3 kcal mol^−1^) contribution and an unfavorable entropic (−TΔS = +23.2 kcal mol^−1^) contribution. Reaction conditions: 20 mM potassium phosphate, 180 mM KCl, pH 7.2, 25 °C. For FRET experiment, [O1] = 200 nM; for POT1 experiments, [POT1] = 920 nM.

Stopped-flow fluorescence experiments to determine the rate of FRET-labeled O1 to POT1 are shown in Figure 2. Binding was observed to be fast, and was complete in about 0.2 s. These data were monophasic and fit to a simple single-exponential reaction model. Surprisingly, the rate of binding was apparently independent of POT1 concentration with an average relaxation time of 80.0 ± 0.4 ms observed over a range of 50 – 600 nM POT1. The apparent lack of any concentration dependence can arise from the existence of a hidden bimolecular interaction step that is faster than what can be observed using our stopped-flow method. If such is the case, the minimal reaction mechanism for O1 binding is

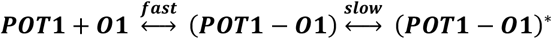

where a fast bimolecular binding step is followed by a slower unimolecular conformational rearrangement of the complex. Only the slower step is resolved in our stopped-flow experiments (Figure 2). Such a mechanism predicts that the rate of the slow second step is constant when the concentration of POT1 is in excess of O1, as is in fact observed in Figure 2. Under such conditions, the observed slow step is characterized by a reciprocal relaxation time of 1/*τ*_2_ = k_2_ + k_−2_, where the rate constants refer to the forward and reverse reactions in the slow step.

**Figure 2.**
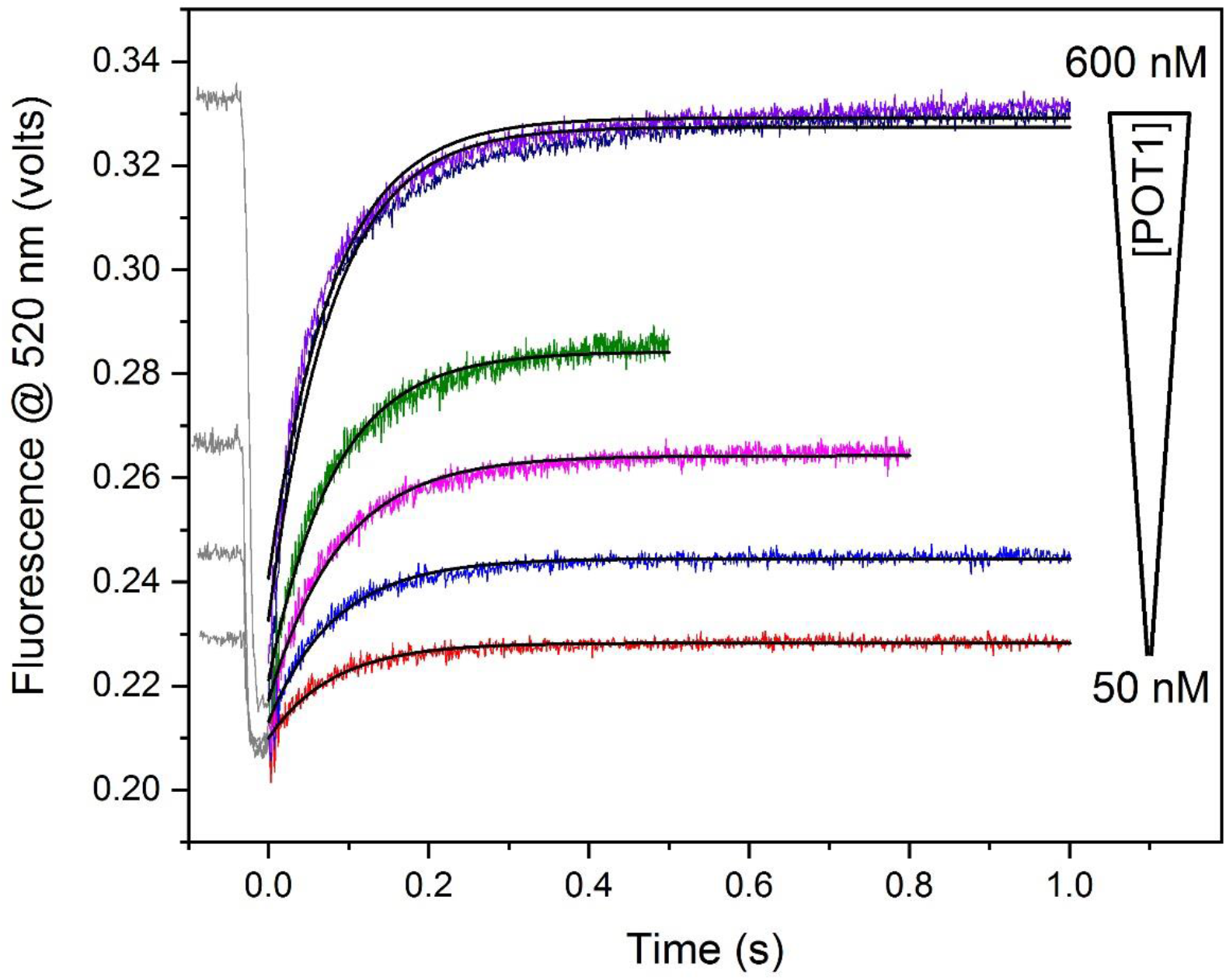
Kinetics of POT1 binding to FRET-labeled O1 determined by stopped-flow mixing experiments. Changes in O1 donor fluorescence are shown as a function of time for increasing concentrations of added POT1. All data were fit to a single exponential relaxation model (solid black lines). The fitted relaxation times were found to be independent of POT1 concentration, with an average value of 82 ± 4 ms. Reaction conditions: The concentration of FRET-O1 was 200 nM in 20 mM potassium phosphate, 180 mM KCl, pH 7.2 with 0, 50, 95, 115, 280, 500 and 600 nM POT1 (concentrations after mixing). The temperature was 25 °C.

### The binding of POT1 to an initially folded telomeric G-quadruplex structure is slow because of conformational selection, the mandatory coupling of binding to a rate-limiting G4 unfolding reaction

Figure 3A shows the kinetics of binding of POT1 to a FRET-labeled telomeric G4 structure that is initially fully folded. The G4 structure is an antiparallel “hybrid” or “3+1” form containing stacked G-quartets, two lateral loops and one side “propeller” loop. In this assay, increased fluorescence results as the G4 structure unfolds and the ends of the quadruplex forming sequence separate. In sharp contrast to the fast binding of single-stranded O1, these data show that POT1 binding to G4 is multiphasic and slow, with an apparent average relaxation time (when the overall reaction is 67% complete) of 2000-3000 s. This is about 25000 times slower than POT1 binding to the single-stranded O1. Three relaxation times of approximately 80 s, 900 s and 16000 s are needed to accurately fit the time course of the reaction. From our previous kinetic studies of telomeric G4 structures (29), we know that G4 unfolding is multiphasic and slow, and is similar to the observed time range of the POT1 binding reaction. To show this directly, we compared the rate of POT1 binding to the rate of G4 unfolding using the complement trap method, in which DNA (or PNA) with a complementary sequence to the G4 strand (O1c, Table S1) is added in excess to drive the rate-limiting G4 unfolding and subsequent fast duplex formation. Figure 3A shows that the rate of POT1 binding tracks almost exactly with the rate of G4 unfolding driven by added complementary DNA or by a smaller γ-PNA sequence. Figure 3B shows a transformation of the data in which the extent of G4 unfolding by complementary trap sequences is compared directly to the extent of unfolding by POT1, showing only slight deviations from a linear correlation. These data emphasize that POT1 and complement trap sequences must act by the same mechanism.

**Figure 3.**
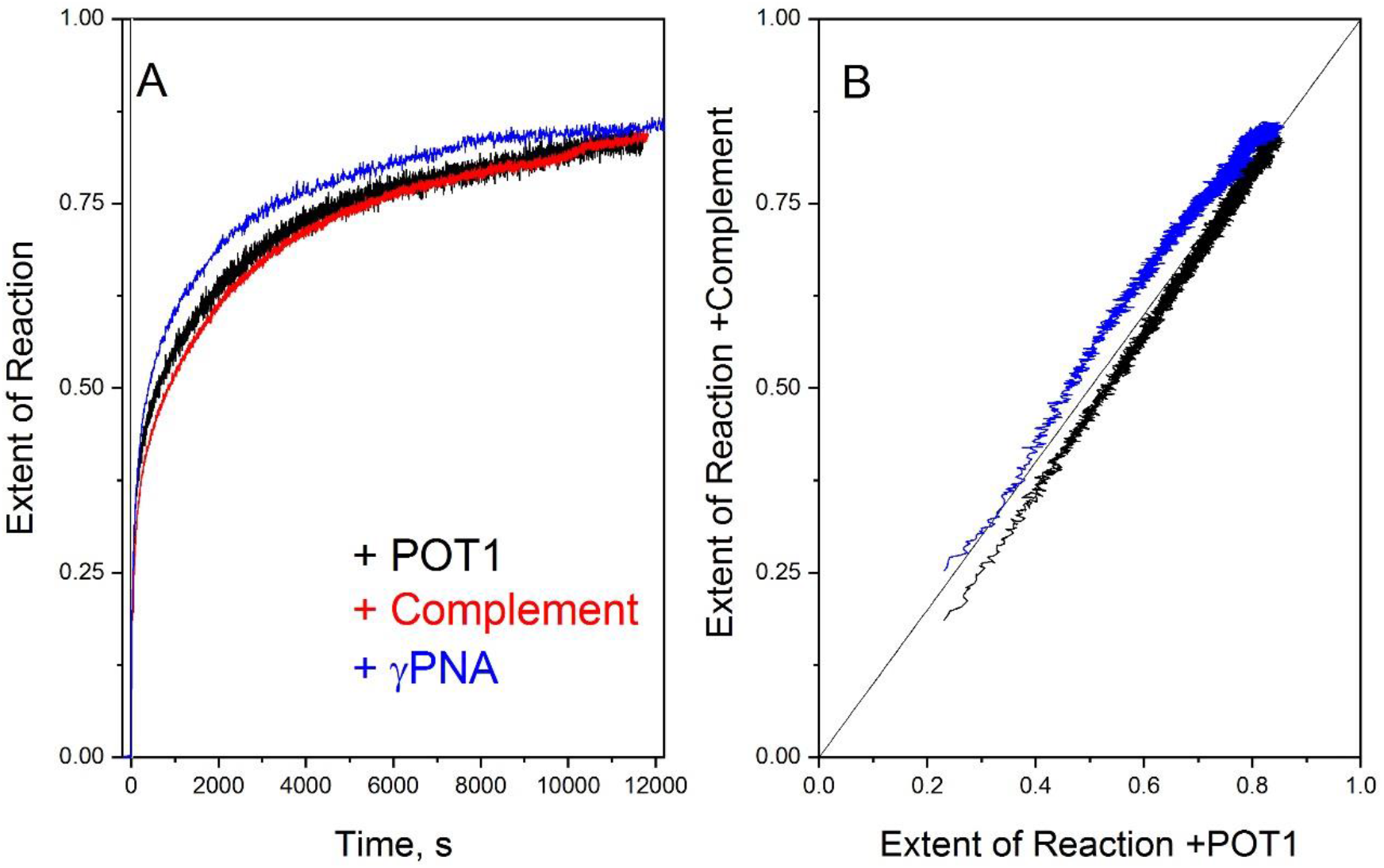
Kinetics of G-quadruplex unfolding by POT1 and DNA or γPNA complementary sequences. **(A)** The unfolding of the FRET-labeled tel22 (d[AGGG(TTAGGG)_3_]) G-quadruplex determined by hand mixing experiments with changes in donor fluorescence monitored as a function of time. The response has been normalized to show the extent of the reaction with respect to endpoint in order to emphasize kinetic similarities. The black line shows unfolding by POT1. The red line shows unfolding using a 22 nt DNA complement to the tel22 sequence as a trapping reagent (29). The blue line shows unfolding by the γPNA H-CCCTAA-NH_2_. There are negligible differences in the time courses of the unfolding reaction for three unfolding agents. **(B)** Plot showing the extent of reaction for the DNA (black) or γPNA (blue) complements with respect to POT1. The diagonal line show the expected correlation if the unfolding reactions had exactly the same kinetics. These data emphasize that the kinetic difference of the nucleic acids with respect to POT1 are negligible. Reaction conditions: 20 mM potassium phosphate,180 mM KCl, pH 7.2, 25 °C, 200 nM Fret-Tel22, 2 μM γPNA, 2 μM complement or 1.8 μM POT1.

These kinetic results strongly suggest that the interaction of POT1 with an initially folded telomeric G4 structure is governed by a conformational selection mechanism in which binding is coupled to mandatory unfolding of the G4 structure. POT1 binding to single-strand DNA is fast, as shown above, so the slow kinetics seen in Figure 3 must result from the slow, rate-limiting unfolding reaction. The overall coupled reaction is

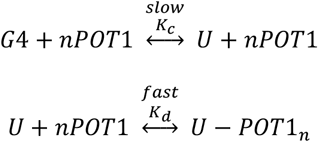

where U represents the unfolded G4 and n is the POT1 binding stoichiometry. The two reactions are coupled through U, the unfolded G4 intermediate.

We also studied the rate of POT1 and DNA complement unfolding of telomeric G4 by circular dichroism (CD) monitored at 295 nm, a wavelength selective for G4 formation (Figure S2). Interestingly, CD showed a simplerunfolding process with two relaxation times of approximately 20 s and 200s, both faster than observed by the FRET measurements shown in Figure 3. The results can be easily reconciled. CD would be highly sensitive to even slight disruption of G-quartet stacking within the G4 structure (for example removal of one strand segment to disrupt the G-quartet structure), whereas FRET is sensitive to the distance between the 5’ and 3’ ends of the quadruplex forming sequence. If binding of multiple POT1 molecules is required to fully unfold the G4 structure (and it is, as will be shown), binding of the first might eliminate the G4-specific CD signal, while additional binding might be needed to fully extend a partially folded strand to yield the maximal FRET change.

### Two POT1 molecules bind to the unfolded 24 nt G4 DNA

We studied the product resulting from POT1 unfolding of G4 by analytical ultracentrifugation (AUC), with the results shown in Figure 4. The results show the distribution of sedimentation coefficients, c(s), at increasing molar ratios of added POT1. From these data the binding stoichiometry can be inferred. Figure 4A shows the sedimentation positions of the folded G4 alone (1.97 S, ≈7600 Da, calculated M_w_ 7575) or of POT1 alone (2.83 S, ≈33000 Da, calculated M_w_ 38860). The vertical black and red lines track these positions in the remaining panels. Figures 4B-E show that with increasing molar ratios of added POT1, the amount of folded G4 decreases as complexes with POT1 form. Initially, at a 0.5:1 POT1:G4 ratio, a species is seen near 4 S that corresponds to a 1:1 complex of POT1:G4 (Figure 4B). As more POT1 is added, the c(s) distributions shift to even higher molecular weight species. Figure 4E shows the ultimate formation of a species with 4.83 S and an apparent molecular weight of ≈75000 Da (dashed horizontal line), as well as some unbound POT1 (red vertical line). The mass of the complex corresponds to the binding of 2 POT1 molecules to the unfolded 24 nt G4 DNA, clearly establishing a binding stoichiometry of 2:1. A separate, two-wavelength, analysis of this same data using the known molar extinction coefficients for the G4 DNA and POT1 provided an estimate of 2.2:1 for the binding stoichiometry.

**Figure 4.**
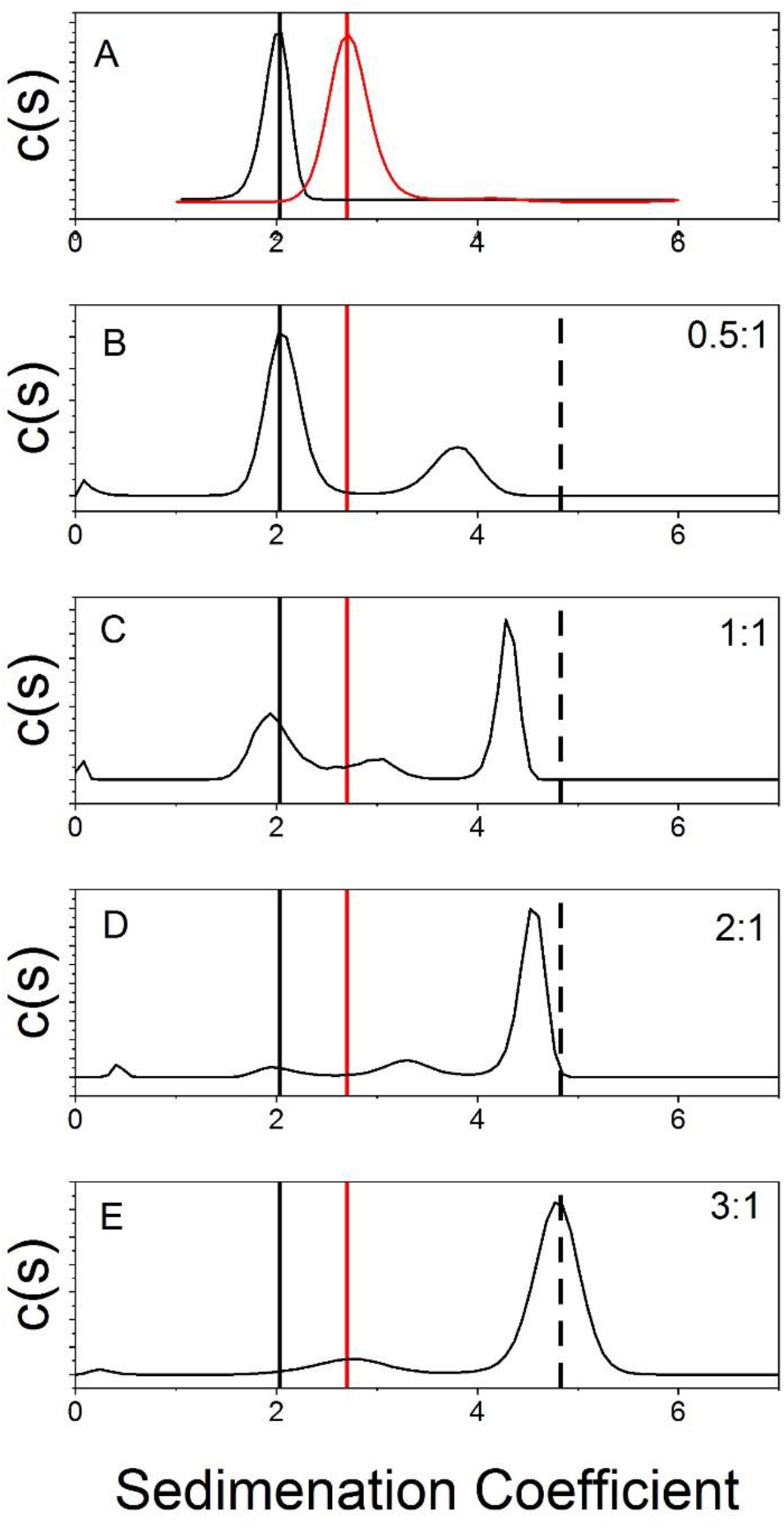
Determination of POT1 binding stoichiometry by analytical ultracentrifugation. The interaction of POT1 with the 2GKU sequence that forms a hybrid 1 G-quadruplex structures was studied. The distribution of sedimentation coefficients (c(s)) is shown for different molar ratios of added POT1. (A) Sedimentation of the 2GKU quadruplex (black) and free POT1 (red) are shown. (B) Sedimentation of a 0.5:1 molar ration of POT1:2GKU. The black and red lines show the position of free 2GKU and POT1, respectively. The new peak near 4S results from the formation of a 1:1 complex. (C)-(E) Sedimentation distributions for increasing molar ratios of POT1:2GKU, as indicated by the labels in the upper right corner of each panel. As the amount of POT1 increases, the amount of 2GKU (near 2S) is depleted and a complex corresponding to a 2:1 molar ratio (shown by the dashed line) is formed. Reaction conditions: Reactions were carried out in POT1 buffer (20 mM KPO_4_, 180 mM KCl, pH 7.2) at room temperature. The 2GKU concentration was 2.5 uM in all experiments while POT1 concentration was varied from 1.25-7.5 μM in B-D. Samples were incubated overnight at room temperature prior to AUC analysis.

### Equilibrium studies of POT1 binding to telomeric G4 by conformational selection

Figure 5 shows the binding isotherm for the interaction of POT1 with an initially folded telomeric hybrid G4 structure. Determination and subsequent analysis of this isotherm was both difficult and time consuming. First, because of the slow kinetics described above, each point on the isotherm was determined independently using a separate solution containing a mixture of 200 nM FRET-labeled G4 and varying concentrations of POT1. Each solution was incubated overnight to ensure that equilibrium was attained before FRET measurements. Second, analysis of the binding isotherm by nonlinear least-squares fitting was also challenging. The isotherm is, by eye, slightly sigmoidal in shape, indicative of a complex binding mechanism. For “simple” binding with one class of sites, a smooth hyperbolic shape would be expected. We explored several binding models to fit the data (shown in Table S3), including “simple” binding, the phenomenological Hill equation and a conformational selection model that explicitly accounts for the coupling of binding to G4 unfolding.

**Figure 5.**
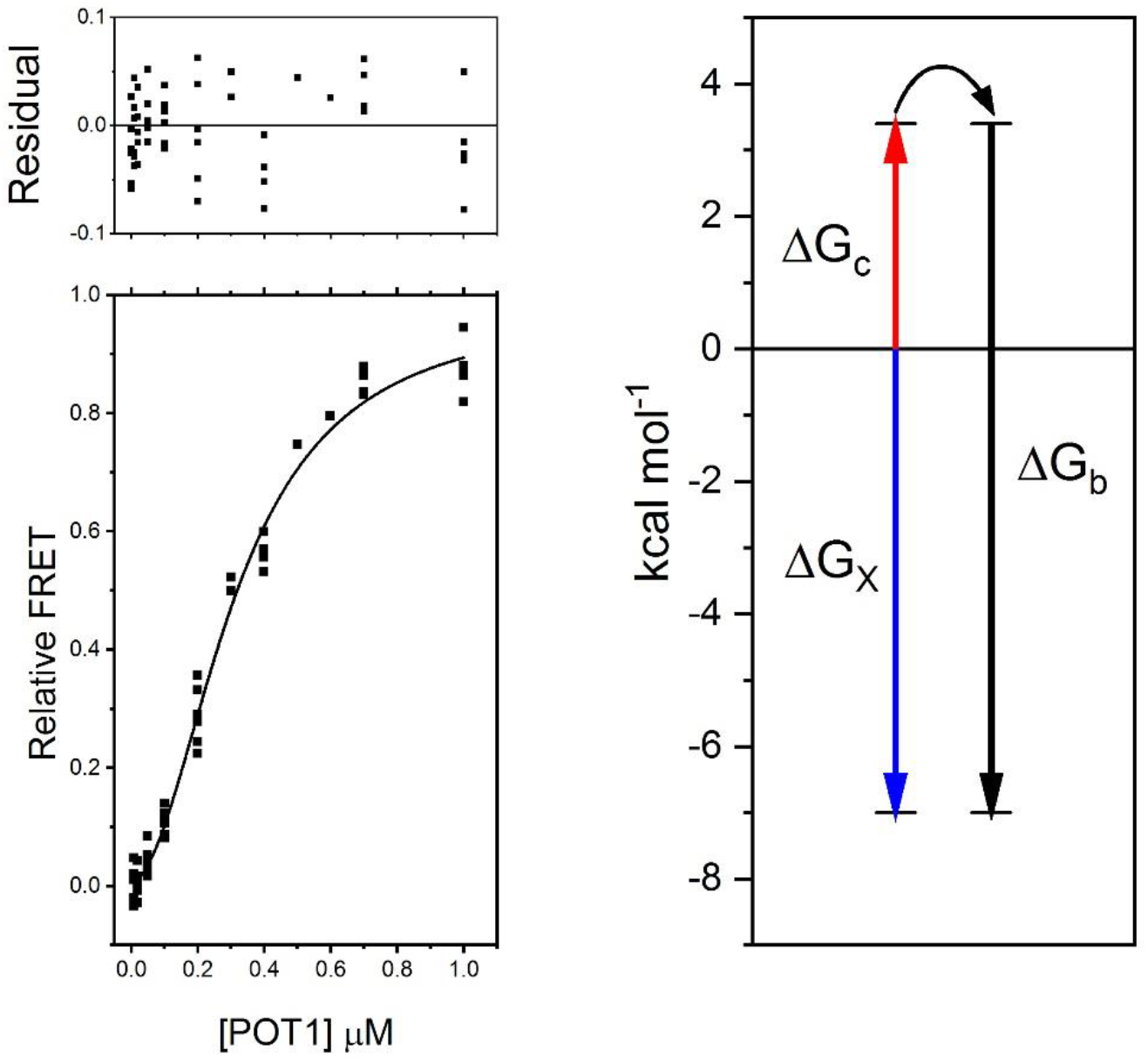
Coupled binding of POT1 to an initially folded G-quadruplex. **(A)** Binding isotherm for the interaction of POT1 to a FRET-labeled G-quadruplex. Each point represents a separate reaction mixture that was allowed to equilibrate for 24 hrs. The solid line represents the best fit to a conformational selection binding model in which POT1 binding is coupled to a mandatory unfolding of the G-quadruplex, as described in the text. **(B)** Free energy diagram illustrating the thermodynamic linkage between POT1 binding (ΔG_b_) and G-quadruplex unfolding (ΔG_c_). The large negative coupling free energy (ΔG_x_) shows that the intrinsic binding affinity of POT1 for single-stranded telomeric DNA drives the unfolding of the G-quadruplex. Reaction conditions: 20 mM potassium phosphate,180 mM KCl, pH 7.2, 25 °C, 200 nM FretG4.

Figure S3 shows residual plots from fits to various models, with the best parameter estimates shown in Table S3. The “simple” model with one class of binding site indicates that approximately 2 POT1 molecules are bound with a K_d_ =0.95 μM. But this model shows nonrandom residuals (Figure S3A), indicating that it fails to account for the sigmoidal character of the binding isotherm. A fit of the data to the Hill model (Figure S3B) with K_d_ =0.32 μM and n=1.9 shows more random residuals, indicative of a better fit. Comparison of the statistics of fits to the two models (Table S3) confirms this, with a clear decrease in the absolute sum of squares of the residuals (ASSR) and the Sy.x parameter that accounts for differences in the number of degrees of freedom resulting from added fitting parameters. Rigorous comparison of the one site and Hill models shows that the choice of the Hill model is statistically warranted, even though it has an additional fitting parameter, with a P value <0.0001. This confirms that the sigmoidal character of the binding isotherm in Figure 5A is significant and needs to be accounted for in any model. While the Hill model adequately describes the data in Figure 5A, it is phenomenological and does not fully describe important details in the underlying reaction mechanism (31,32). In particular, for the POT1-G4 interaction, the Hill model neglects the linkage of G4 unfolding to the binding process and fails to account for the energetic contribution of the unfolding step.

We therefore considered a more physically meaningful binding model, the simplest of which is a restricted conformational selection (CS) model as described above. The fitting function for the restricted conformational selection model is

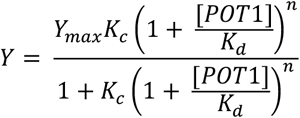

where K_c_ is the equilibrium constant for G4 unfolding, K_d_ is the dissociation constant for POT1 binding to the unfolded strand, and n is the binding stoichiometry. A detailed discussion of the model and the fitting function was presented in previous work from our laboratory (33).

Figure 5 shows the successful unconstrained fit of the binding data to the restricted conformational selection model, with K_c_=0.003, K_d_=0.02 μM and n=2.1 (Table S3). This model fits the data as well as the Hill model albeit with more fitting parameters (Table S3). While the choice of the CS model might be criticized on purely statistical grounds, the fact that it provides a physically more meaningful understanding of the underlying reaction mechanism leads us to favor it over the Hill model. The fitting of the data to the model was challenging, primarily because of the high correlation of parameter estimates. We did extensive housekeeping to explore the quality of the fit. First, Figure S4 shows the somewhat unusual error spaces of parameter estimates. These map the reliability of parameter estimates and show unexpected asymmetry. For both K_c_ and K_d_, there is a broad minimum in the error space. Second, we did Monte Carlo simulations (not shown) of our best fit, the results of which confirmed the correlation of parameter estimates and rather broad distributions for quantitative estimates of both K_c_ and K_d_.

As a reality check of the fitted parameters, we compared the results of the fit to independently determined values as shown in Table 1. We previously studied in detail the thermodynamics of G4 unfolding by thermal denaturation (22). For the first two steps of a multistep unfolding process, we measured a free energy change of +4.0 kcal mol^−1^ at 25 °C, which converts to an unfolding equilibrium constant of 0.001, remarkably close to our fitted value of 0.003. The average dissociation constant for the interaction POT1 with the single-stranded O1 was 0.038 μM (Figure 1, Table S2), close to the fitted value of 0.02 μM. Finally, the stoichiometry of POT1 binding was determined to be near 2:1 by fitting our data, in excellent agreement with the value determined independently by AUC. As an alternate fitting strategy, we used each of these independently determined parameters in constrained fits with the results shown in Table S3. This strategy alleviates problems with the correlation of parameter estimates, and yields similar estimates for the unconstrained parameters compared to those obtained in unconstrained fits. Overall, the physically meaningful conformational selection model provides a reasonable description of the binding isotherm in Figure 5, and yields meaningful quantitative estimates for the underlying equilibrium constants for the steps in the reaction. These quantities can be used to construct the free energy diagram shown in Figure 5B that concisely illustrates how POT1 unfolds a G4 structure. The diagram shows that the energetic cost of unfolding is overcome by the favorable free energy of POT1 binding to the exposed single-strand. Figure 5B explicitly defines the thermodynamic linkage between POT1 binding and G4 unfolding.

**Table 1.** Comparison of fitted conformation selection parameters to independently determined values.

| Parameter | Fit Value | Independent Estimate | Comment |
| --- | --- | --- | --- |
| $K_c$ | 0.003 | 0.0011 | First steps in G4 unfolding determined by thermal denaturation experiments (22) |
| $K_d$ μM | 0.02 | 0.038 | Binding of O1 to POT1 (Figure 1, Figure S1, Table S1) |
| $n$ | 2.1 | 2.0 | Determined by AUC (Figure 4) |

### POT1 unfolds all G4 forms of telomeric DNA, but not duplex DNA or a promoter G4 structure

Human telomeric DNA sequence can adopt several G4 structural forms depending on the exact nucleotides at the 3’ and 5’ ends of the repeat sequence (Table S1) and the solution conditions, most critically the identity of the monovalent cation present. These structures include an antiparallel “basket”, two different antiparallel “hybrid” structures and a parallel “propeller” structure (Figure 6). We designed experiments to examine how POT1 interacted with these different G4 structural forms, or with duplex DNA or a G4 structure formed by a non-telomeric sequence. Figure 6A shows kinetic FRET unfolding experiments. The data show that the rates of POT1 unfolding of both “hybrid” forms and of the parallel “propeller” form are very similar. In contrast, the rate of unfolding of the antiparallel “basket” form is much faster. In all of these cases, the rate of G4 unfolding by POT1 was essentially the same as the rate of unfolding by a DNA complement trap sequence (data not shown). POT1 unfolds all of these G4 structural forms, but the rate of unfolding depends solely on the intrinsic unfolding rate of the particular structure. The “basket” form in Na+ solution is kinetically less stable than the “hybrid” and “propeller” forms and unfolds at a faster rate (23), accounting for the difference seen in Figure 6A.

**Figure 6.**
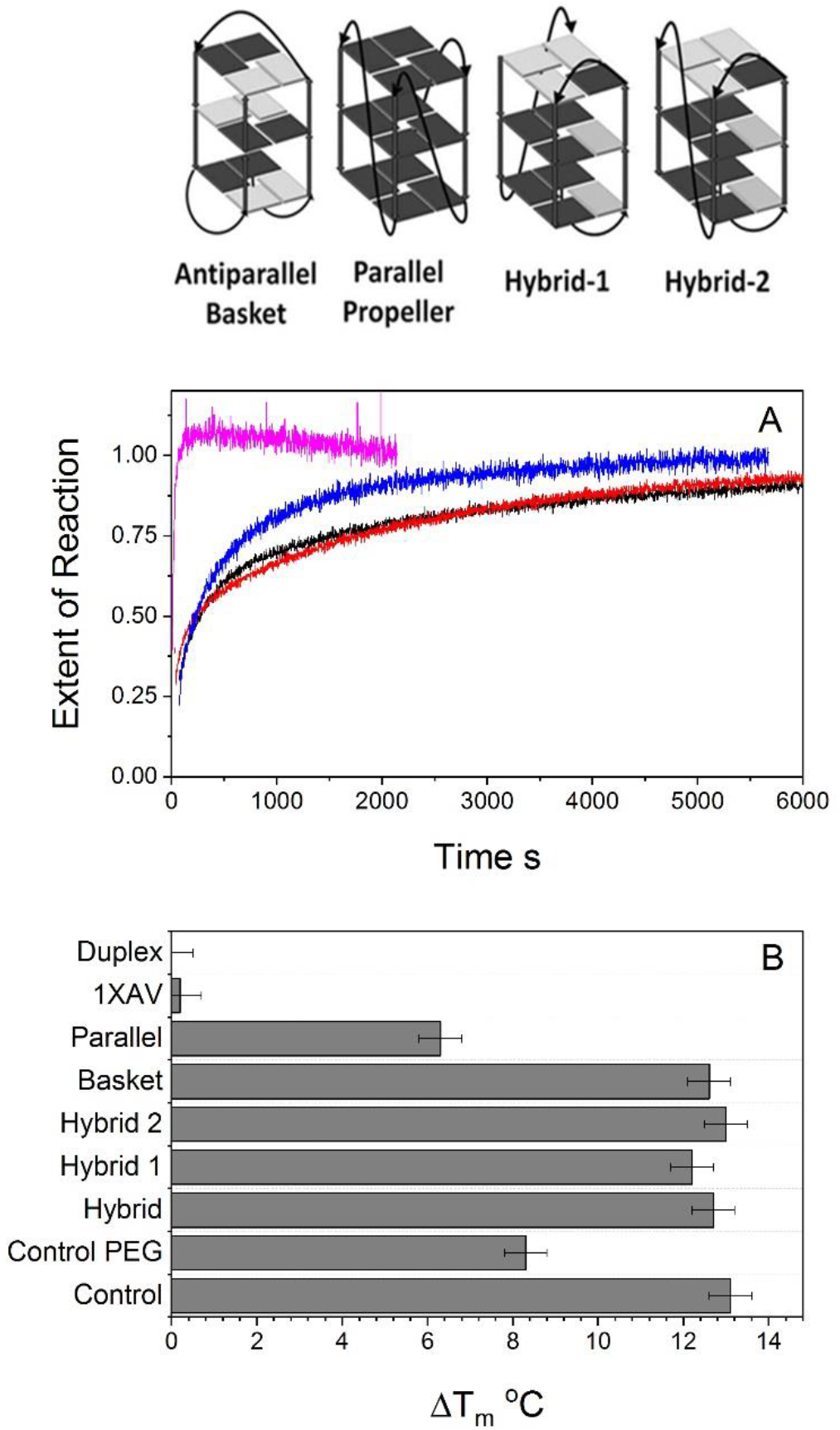
Specificity of POT1 unfolding of G-quadruplex structures. Four structural forms of human telomere G-quadruplexes are shown at the top of the figure. Formation of each structure depends on solution conditions and the exact oligonucleotide sequence. (A) Kinetics of POT1 unfolding of different G-quadruplex forms. The black and red curves show the unfolding of hybrid 1 and hybrid 2 structures, respectively. The blue curve shows unfolding of the parallel propeller form. The magenta curve shows the unfolding of the antiparallel basket form. (B) Results from a differential scanning fluorometry assay for POT1 binding. The shift in the thermal denaturation temperature of POT1 (ΔT_m_) is correlated with the binding affinity for a given nucleic acid structure. The control shows binding to O1, the preferred 10 nt binding sequence. “Control PEG” shows the effect of added 40% polyethylene glycol on the control binding reaction. The data show that all hybrid and basket forms have roughly the same binding affinity for POT1, while the parallel form is slightly less affine. Notably, POT1 does not unfold and bind to a nontelomeric parallel G-quadruplex, 1XAV. Nor does it unfold or bind to a duplex DNA form with the human telomere sequence. (The schematic of telomeric G-quadruplex structures was adapted, with permission, from R. Hansel et al (2011) *Nucl. Acids Res*. 39:5766-75). Reaction conditions for unfolding: 20 mM potassium phosphate,180 mM KCl, pH 7.2, 25 °C, 200 nM FretG4, 1.8 μM POT1.

We also used a differential scanning fluorometry (DSF) thermal shift assay to probe the specificity of POT1 interactions (Figure 6B). This assay is fully described in a companion article in this issue, but will be briefly described here. The assay uses the fluorophore Sypro Orange to monitor the thermal denaturation of POT1 alone or in the presence of DNA binding sequences. DNA binding stabilizes POT1, increasing the temperature at which it denatures (T_m_). POT1 alone, in the DSF assay, has a T_m_ = 51±0.5 °C. The increase in T_m_ upon DNA binding is proportional to the affinity of the binding interaction (along with binding stoichiometry, enthalpy and other factors). As seen in Figure 6B, binding of the control oligonucleotide O1 leads to an increase in T_m_ of 12-13°C. Binding of telomeric G4 structures initially in the “basket” and “hybrid” forms produces similar increases in T_m_, indicating binding similar to the control O1 oligonucleotide sequence. It is important to distinguish here that the DSF assay is a thermodynamic measure of the stability of POT1-DNA complexes, and has no direct connection with the kinetic assay shown in panel A, since samples were fully equilibrated before the start of the assay. DSF also shows that POT1 interacts with the telomeric parallel “propeller” G4 structure (Figure 6B). In this case, there is a slightly confounding issue arising from the conditions necessary for the formation of the “propeller” structure. The “propeller” structure, while seen in crystals, is not the predominant form in solution (34), so it is necessary to add high concentrations of cosolvents to drive its formation. Polyethylene glycol (PEG) is one such cosolvent. We found in a control experiment that addition of PEG decreases the magnitude of the T_m_ shift resulting from O1 binding to 8-9°C (Figure 6B, “Control PEG”). The thermal shift observed for the parallel “propeller” structure in PEG is similar to that value. While at first glance it might appear that the POT1 interaction with the parallel form is weaker than for the basket and hybrid form, that is probably not the case and the apparent difference is instead due to the presence of PEG. These results from DSF indicate that POT1 has little or no preference for any telomeric G4 structural form, and will unfold and bind to all of them equally well.

In contrast, Figure 6B shows that POT1 will not interact with a DNA duplex formed from the telomeric G4-forming sequence and its complementary strand. Nor will POT1 unfold a parallel G4 structure formed by a nontelomeric sequence (Figure 6B, “1XAV”). 1XAV is a G4 structure formed by a modified sequence from the c-myc promoter region, d[TGAG_3_TG_3_TAG_3_TG_3_TA_2_]. These two examples suggest that POT1 acts selectively to stabilize unfolded telomeric G4 structures, a consequence of its selective binding to the single-stranded telomeric sequence. The telomeric duplex is not disrupted presumably because the energetic cost of separating the two strands cannot be overcome by the energy from POT1 binding.

### The POT1-DNA complex is hydrodynamically compact

In all of the G4-POT1 kinetic studies shown above, we normalized the FRET amplitudes of the time courses to emphasize the similarity of the G4 unfolding rates observed for POT1 and for complement trap sequences. In fact, the overall changes in FRET amplitudes observed at the end of the unfolding reactions are different (Figure 7A). Figure 7A shows that the amplitude of the FRET donor fluorescence peak at 520 nm is 4.5-fold larger for the complement-trap complex than for the POT1 complex. This difference means that the separation of the donor and acceptor are different for the two complexes, so their shapes must be different. Since FRET depends on the sixth power of the distance separating the donor and acceptor, Figure 7A indicates that the ends of the G4 sequence are farther apart in the duplex formed by addition of the complementary trap sequence than they are in the POT1 complex. For the complementary trap complex, the G4 unfolds and then forms a rod-like duplex molecule with the ends of the G4 strand maximally separated. In contrast, the POT1 complex must have a less extended shape with the ends of the single-stranded G4 sequence separated to a lesser extent. The spectra in Figure 7A can, in principle, be used to estimate the end-to-end distances of the labeled G4 sequence within the complexes, although with difficulty because of numerous assumptions needed and because the of the extended linkers used to attach the probes to each end. We nonetheless tried, with our best estimates showing that the donor and acceptor are separated by 74 ± 8 angstroms in the duplex, but only by 62 ± 5 angstroms in the POT1 complex. For reference, in the folded G4 structure, the donor-acceptor distance was calculated to be 46 ± 4 angstroms. We will soon show molecular models that can explain these distances.

**Figure 7.**
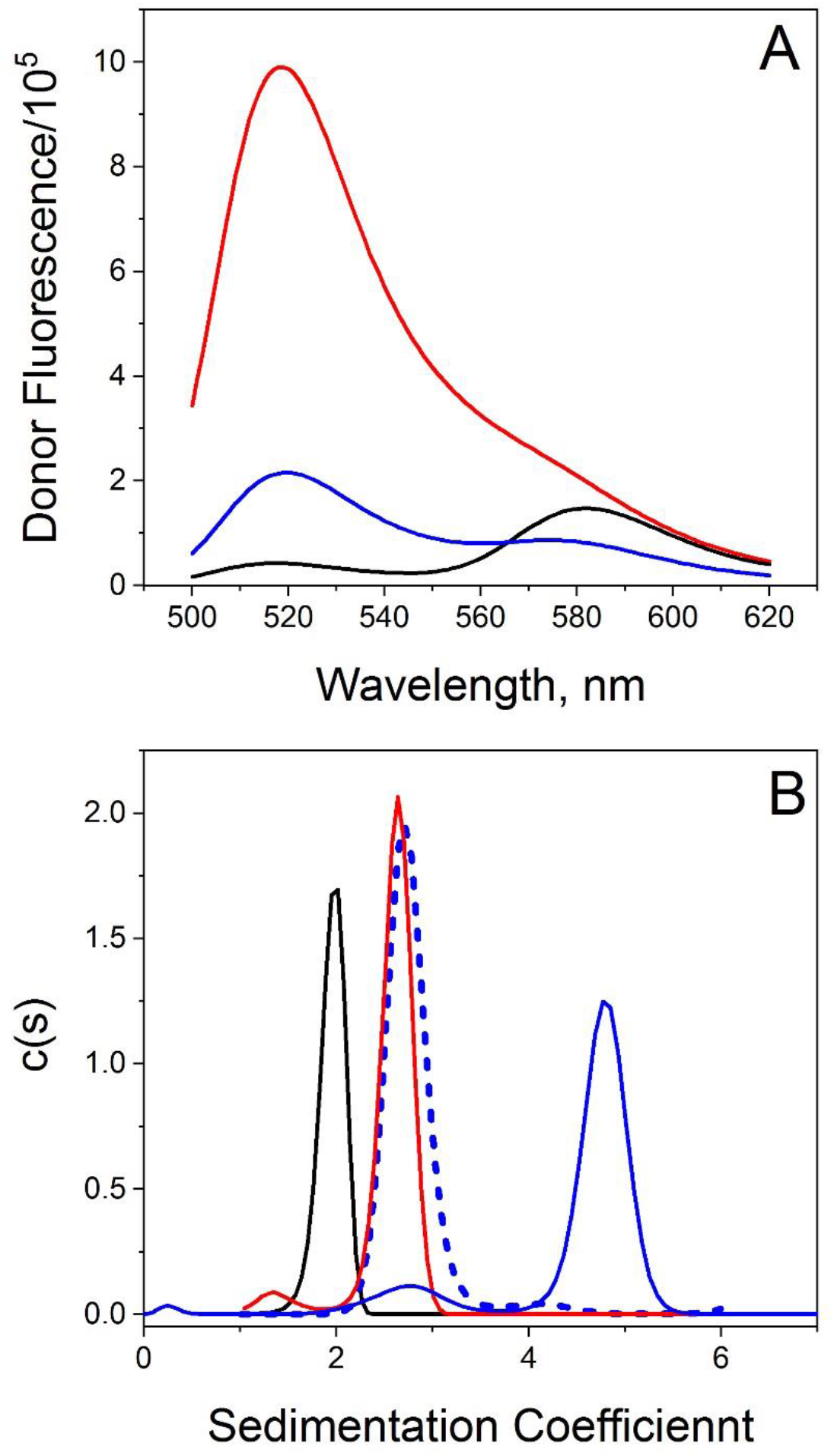
Characterization of the POT1-DNA complex. (A) Excitation spectra of folded and unfolded FRET-labeled 2GKU. The black curve shows the folded G-quadruplex form. The blue curve shows the 2:1 POT1-2GKU complex. The red curve shows the duplex form of the 2GKU sequence produced by addition of its complementary strand. The differences in the donor fluorescence peak near 520 nm signify differences in FRET efficiency that is proportional to the end-to-end distance of the 2GKU strand. (B) Analytical ultracentrifugation of 2GKU forms. Sedimentation coefficient distributions (c(s)) are shown. The black curve shows the folded G-quadruplex form. The dashed blue curve shows free POT1. The blue curve shows the 2:1 POT1-2GKU complex. The red curve shows the duplex form of 2GKU.

Figure 7B and Table 2 show AUC results that further characterize the hydrodynamic shapes of POT1 and complement trap complexes. The plots show sedimentation coefficient distributions (c(s)), from which frictional ratios (f/f_0_) were estimated for each sample. The frictional ratio is defined as the ratio of the frictional coefficient experienced by the sedimenting molecule relative to that of an ideal sphere with the same molecular weight. The value of f/f_0_ is a measure of hydrodynamic shape, with spherical molecules having a ratio of 1.0, and more asymmetric molecules having larger ratios. For non-spherical prolate or oblate ellipsoids, the frictional ratio depends on the exact dimensions of the major and minor axes. The value of f/f_0_ depends on shape and is independent of molecular weight. We found that the duplex complex has S_20,w_ = 2.63±0.05 and f/f_0_=1.58, consistent with a rod-like molecule. The POT1-complex sediments faster, with S_20,w_ = 4.85±0.05 and f/f_0_=1.15, indicative of a more compact structure. For frictional ratios less than 1.2, it is difficult to distinguish prolate from oblate ellipsoid shapes. These data are qualitatively consistent with results of the FRET measurements above.

**Table 2.**
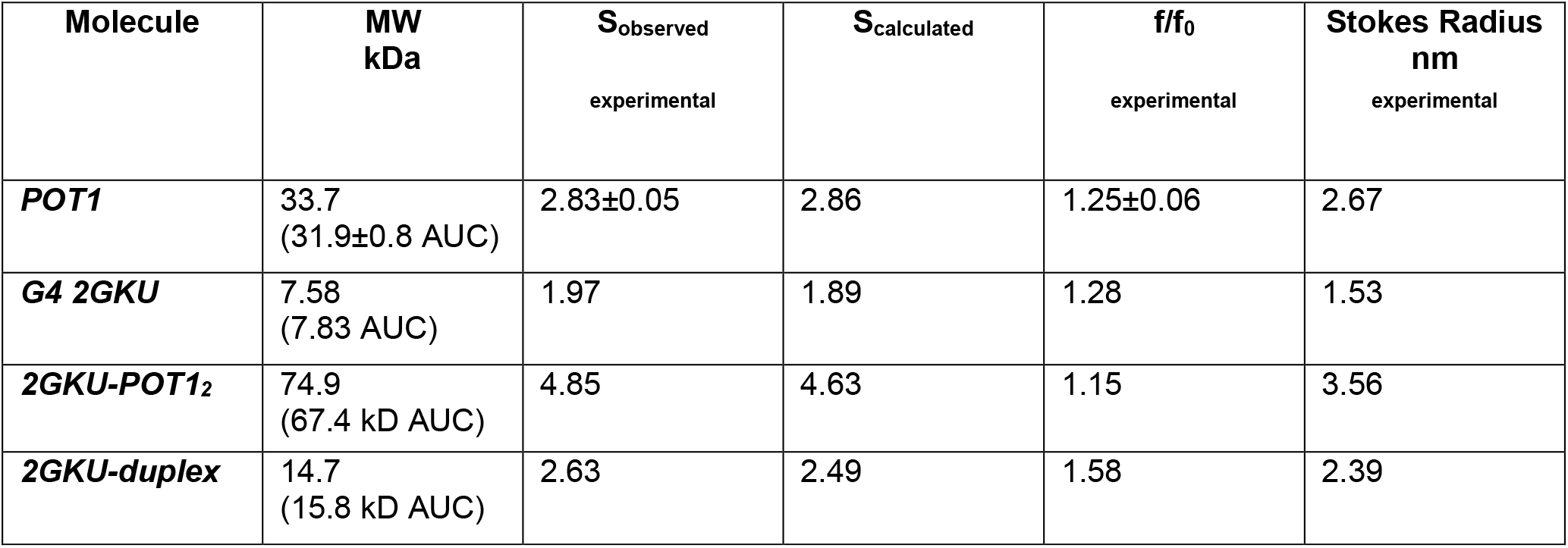
Hydrodynamic properties of G4 complexes.

Figure 8 shows the results of molecular dynamics simulations that provide plausible molecular models for the duplex and POT1 complexes. Addition of the complement trap sequence results in the formation of a canonical duplex complex as shown in Figure 8A. Our model predicts a sedimentation coefficient of 2.86 S, in excellent agreement with the experimentally determined value (Table 2). The model of POT1-DNA complex (Figure 8B) is more complicated, with two POT1 molecules bound to the single-stranded DNA that bends to form a crescent-like shape. Figure S5 shows the POT1 complex in different orientations. Both POT1 molecules, consisting of OBD 1 and 2, form a continuous basic concave groove where the optimal DNA binding sequence lies, inherently bending the DNA strand. This reduces the distance between the 5’- and 3’-bases where the FRET labels are attached. Our model predicts a sedimentation coefficient of 4.63 S, an estimate within 5% of the experimentally determined value (Table 2). The 5’-OH and 3’-OH oxygen atoms were used for end-to-end distances for the DNA in both complexes were measured over the entire 10 ns trajectory of the simulation (Figure S6). For the duplex, the end-to-end distance is 78.9±2.9 Å, in good agreement with the 74 ± 8 Å experimental estimate obtained by FRET. In the POT1 complex model, the distance is 55.2 ±3.5 Å, compared to the experimental estimate of 62 ± 5 Å. The values for the models and experiments are within 6-11% of one another, suggesting that the models are reasonable representations of the complexes that form. It should be noted that the molecular dynamics simulations did not contain the actual FRET labels, so it is the trend in the distribution of distances that is important, not the absolute distances.

**Figure 8.**
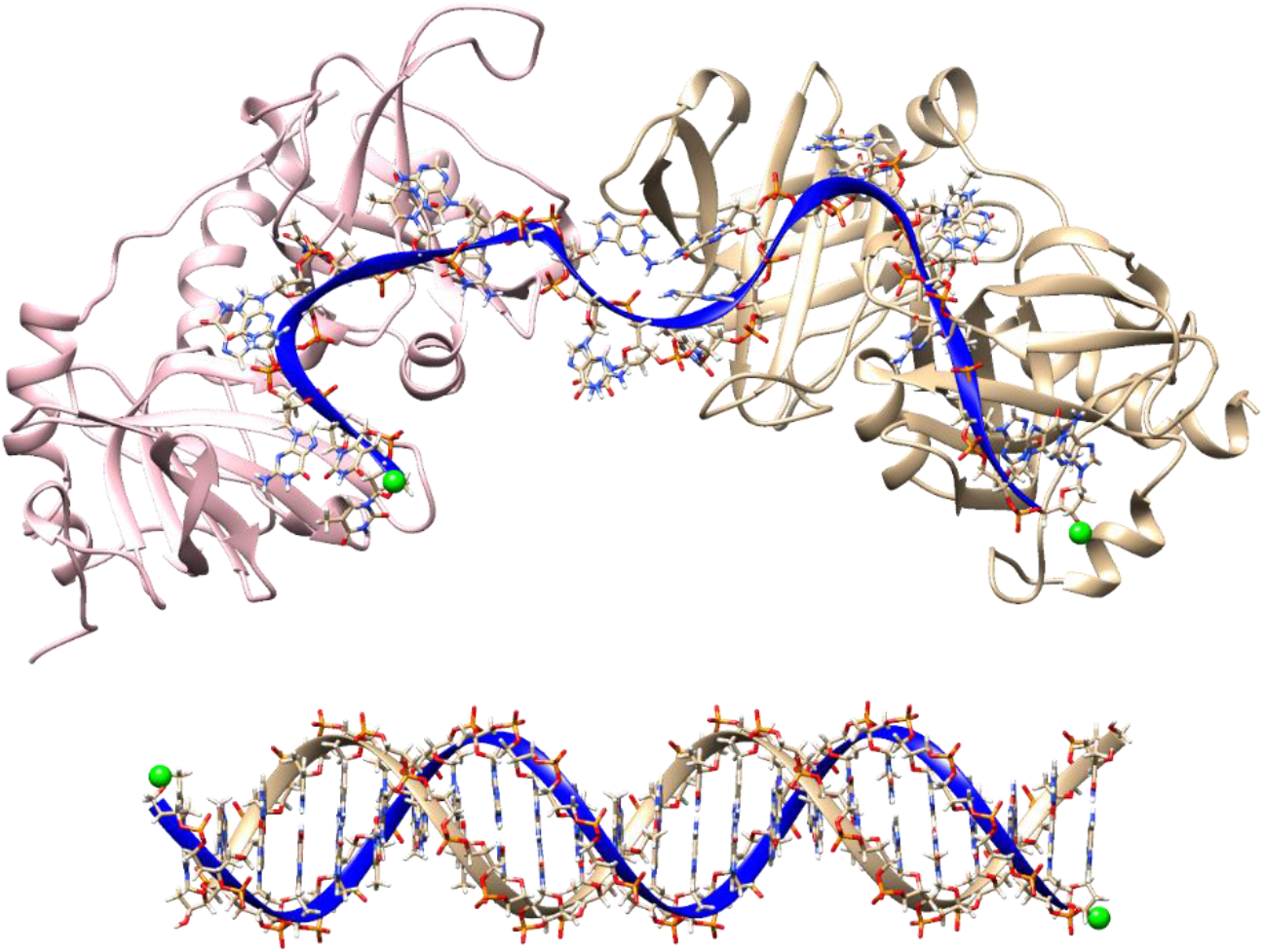
Structural model of the POT1-DNA complex. (Top) MD-derived 2GKU-POT1_2_ complex showing two POT1 proteins (ribbons shown in pink and tan) bound to the single-stranded 2GKU oligonucleotide (ribbon shown in blue). For each POT1 protein the ssDNA spans both OB1 and OB2, binding in the continuous basic concave groove, consistent with the crystal structure 1XJV. Binding of the DNA to the POT1 OB1 and OB2 domains leads to an approximately 90° bend in the DNA backbone. There is free rotation about the connecting DNA sequence in between the two POT1 proteins, which leads to an overall ‘w’ shape (although it can twist). Green spheres located at the 3’ and 5’ ends of either structure highlight the oxygen atoms used in distance calculations.

## Discussion

These studies show that POT1 binds to telomeric G4 structures by a conformational selection mechanism in which binding to the single-stranded repeat sequence is preceded by an obligatory unfolding of the quadruplex. We found no evidence for any interaction between POT1 and a folded G4 structure. Our data and analysis provide quantitative characterization of the underlying reaction. The overall free energy of POT1 unfolding of, and binding to, telomeric G4 (*ΔG_Total_*) is a sum of two contributions

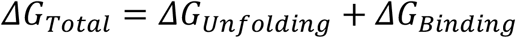

where *ΔG_Unfolding_* and *ΔG_Binding_* are the free energies for G4 unfolding and POT1 binding, respectively. Figure 5B shows how the +3.4 kcal mol^−1^ unfavorable free energy cost to unfold G4 is overcome by −10.5 kcal mol^−1^ binding reaction energy to yield a favorable overall reaction of −7.1 kcal mol^−1^. This conformation selection model is consistent with previous suggestions (15,19,20) that POT1 might “trap” an unfolded G4 intermediate but is inconsistent with suggestions that POT1 interacts initially with intact G4 structures or nearby sequences to destabilize, then unfold, telomeric quadruplexes (16,20).

We previously characterized the complement unfolding reaction in some detail (23,29) as essential background for understanding POT1 unfolding of G4. The kinetics of telomeric G4 unfolding by POT1 track with the unfolding reaction triggered by the addition complementary DNA or PNA sequences, as shown in Figures 3 and 6. These kinetics are fully consistent with the conformational selection model, and show that the rate limiting step in POT1 binding to initially folded G4 structures is the obligatory unfolding process. We show that binding of POT1 to its single-stranded binding site is fast (Figure 2). Our finding that complementary DNA or PNA sequences can mimic POT1 unfolding of G4 structures is consistent with the early report from the Cech laboratory (15), but is inconsistent with a subsequent atomic force microscopic study (16). Direct binding of POT1 to a folded G4 structure would be expected to be a fast bimolecular interaction on the same time scale as we observed for the POT1-O1 reaction. We found no evidence for such an interaction.

We find that POT1 can unfold all telomeric G4 conformational forms, with the rate of unfolding tracking with the intrinsic unfolding rate for each form. The slow kinetics we observe for the G4-POT1 interaction may, at first glance, raise questions about the “physiological” relevance of the reaction. However, the complete cell cycle of human cell lines typically takes about 24 hours to complete (BNID112260,(35)) and more specifically telomere extension in human fibroblast cells was shown to take >10 hours during the S phase (36). The rates we observe for POT1 reactions are compatible with the times of these *in vivo* processes.

FRET measurements and hydrodynamic studies show that the POT1-ssDNA complex is a compact assembly, relative to a DNA duplex structure formed by the same sequence. A stable atomistic model (Figure 8, Figure S5) provides details of the assembly, and shows two POT1 molecules bound to the 22 nt telomeric DNA sequence. The POT1 molecules are each bound through their two N-terminal OB domains, as was described in the crystal structure from the Cech laboratory (10). This atomic model accurately predicts measurable experimental hydrodynamic properties, with the sedimentation coefficient prediction within 4-5% of the observed value (Table 2). Our model seems fully consistent with the electron microscopy studies of POT1 bound to long telomeric sequences (13), which showed ordered, compact globular assemblies.

In humans, the 3’ single-strand telomeric overhang is 200±75 nt long (37) and can form multiple G4 units. Our present results provide a firm basis for future studies of POT1 interactions with such longer, perhaps more natural sequences. These studies are underway in our laboratory. We have carefully characterized the structures of long telomeric sequences (17,38–40) by molecular dynamics simulations and experimental biophysical studies and have shown that for lengths up to 196 nt they always fold into structures containing the maximal number of possible G4 units, with no single-stranded gaps longer than 3 nt. POT1 would thus encounter an ordered array of G4 structures as a binding substrate in these longer sequences. Our preliminary kinetic and equilibrium binding studies (not shown) indicate that a conformational selection mechanism similar to that described here is also operative for POT1 binding to long telomeric sequences.

## SUPPLEMENTARY DATA

Supplementary Data containing Material and Methods, Tables S1–S3 and Figures S1–S6 are available at NAR online.

## Supplementary Data

### MATERIALS AND METHODS

#### POT1 Protein Expression and Purification

A plasmid encoding an N-terminal transcript variant 2 of human POT1 (POT1-N used to determine the crystal structure 1XJV was a generous gift of the Cech laboratory). The POT1-N, POT1 throughout the text, coding sequence was excised from the plasmid and cloned into the pET21a expression vector designed to produce a protein with C-terminal 6His tag. After verifying the coding sequence, the protein was expressed in E. coli strain C41. The biomass was produced in rich (LB) medium in the presence of 100 μg/ml ampicillin at 37° C. Briefly, cultures were inoculated at an optical density A600 nm 0.1-0.15 and grown until reached density of 0.6-0.7 with shaking. Cells were harvested by centrifugation at 5,000 x g for 15 min at 4° C. Pellets were washed with sterile basal M9 medium without additives and harvested again as before. For POT1 production, cells were suspended in M9 medium supplemented with 2% D-glucose, 10 ml/l of Basal Vitamins Eagle medium, 2 mM MgSO4, 0.1 mM CaCl2, and 100 μM FeSO4. Cultures were allowed to adapt to lower growth temperature (18° C) for one hour with shaking in the incubator shaker. Expression of POT1 was induced by addition of IPTG to a final concentration 0.25 mM. After 16-18 hours cells were collected by centrifugation and either directly subjected to protein extraction or kept at −80° C. Cell pellet obtained from 3L of synthetic medium was suspended in 100 ml of lysis buffer containing 50 mM Tris-HCl, pH 7.2, 300 mM NaCl, 10% sucrose 10 mM imidazole, 0.1% NP-40, EDTA-free protease inhibitor cocktail, 1 mM PMSF, 2 mM β-mercaptoethanol. The suspension was sonicated once for 30 seconds (2 sec on/2 sec off) on ice to facilitate formation of homogeneous suspension. A few milligrams of lysozyme was added directly to the suspension and incubated at 4°C for 20 min on a nutator mixer. The extract was adjusted to 10 mM MgCl_2_ and incubated at room temperature on a nutator mixer for 20 min after addition of 400 U of DNase I and 2 mg of RNase A, followed by sonication on ice for 30 sec (2 sec on/2 sec off) a couple of times. We tested the efficiency of DNA digest/shearing by pipetting the extract up and down using yellow tip until no “strings” were detected. It is important to note that high viscosity of the extract will negatively affect following steps of purification. Cell debris was removed after centrifugation at 75,000 xg for 40 min. The POT1 was purified by immobilized metal affinity (IMAC) and anionic exchange chromatography. Briefly, we used an automated Profinia system (Bio-Rad) that allows sequential affinity chromatography and desalting of the sample with Bio-Rad cartridges, Profinity Ni-charged IMAC and P6, respectively. Buffers, as 1X, were used in sequential order: A, 300 mM NaCl, 50 mM Tris-HCl, pH 7.2, 10 mM imidazole, 2 mM β-mercaptoethanol, 10% sucrose; B, the same as A, but 20 mM imidazole; C, the same as A, but 250 mM imidazole; D, (desalting buffer) 150 mM NaCl, 50 mM Tris-HCl, pH 7.2, 5 mM β-mercaptoethanol, 5% sucrose. The desalted sample was concentrated with Amicon centrifugal devices (10 K cut) and loaded on anionic exchange column HiTrap Q HP equilibrated in buffer D, using AKTA Purifier system. The vast majority of POT1 did not bind the resin and eluted in the flow-through fraction. The contaminating proteins bind to the resin and were eluted with increasing concentrations of NaCl. This procedure yields in 90%-95% pure protein, as judged by SDS-PAGE after staining with Coomassie Blue. The purified protein was confirmed to be intact POT1-N by western blot analysis using HIS6-tag monoclonal antibody (H8), mass spectrometry, analytical ultracentrifugation (AUC), and by specific binding to the O1 oligonucleotide d[TAGGGTTAG] (Supplemental Table S1). The purified protein was stored in aliquots of ~50 μM at −80° C in buffer D. Before use, the thawed preparations were transferred into POT1 buffer (20 mM potassium phosphate, 180 mM KCl, pH 7.2) by gel filtration using a BioGel P6 spin column. POT1 concentrations were estimated from the absorbance at 280 nm using a calculated extinction coefficient of 41.37 mM^−1^ cm^−1^

#### Oligonucleotide preparation

Oligonucleotides and their extinction coefficients are listed in Table S1. Unlabelled oligonucleotides were obtained in a desalted, lyophilized state from either IDT (Coralville, IA) or Eurofins Genomics (Louisville, KY). Stock solutions were prepared by adding deionized H_2_O to give ~1 mM concentration and stored at 4 °C. Fluorescently tagged oligonucleotides containing a 5’-6FAM FRET donor and a 3’-Tamra acceptor were synthesized and HPLC-purified by Eurofins Genomics. They were reconstituted in deionized H_2_O to ~100 μM and stored in the dark at 4 °C. Working solutions of oligonucleotides were prepared diluting the stock solution to the desired final concentration in POT1 buffer, denaturing in a 1-L boiling water bath for ~10 min followed by annealing by slow cooling to room temperature. Folding of G4 samples was checked by measuring their CD spectrum or, for the FRET-labelled oligonucleotides, by comparing the emission spectra of unfolded and folded samples determined by exciting 6Fam (495 nm) or Tamra (560 nm).

#### Differential scanning fluorimetry (DSF)

DSF experiments were carried out using an Applied Biosystems StepOne Plus real-time PCR system. Melting curves were determined in 96 well plates using a temperature ramp from 20°C to 99°C. SYPRO Orange dye (1), which preferentially binds to denatured proteins and becomes fluorescent, was used to monitor thermal unfolding of POT1 in the absence and presence of DNA. DNA concentrations were typically at a ten-fold excess to protein (5 μM) in POT1 buffer. Twenty μL of sample was loaded in each well, which was sealed and centrifuged at 1400 rpm for 2 min. Each unfolding reaction was run in duplicate or triplicate and repeated with at least two different plates. POT1 T_m_ was determined from the maximum in the first derivative of the melting curve and ΔT_m_ was estimated from the difference in T_m_ values with and without DNA.

#### Equilibrium Titrations

##### O1 DNA

The extent of POT1-ssDNA binding was determined at 25 °C by measuring changes in fluorescence accompanying serial additions of ligand to receptor using a FluoroMax-3 fluorometer (Jobin Yvon Inc., Edison, NJ). For experiments in which FRET-labelled O1 was titrated with POT1, the labelled DNA was excited at 495 nm and emission measured at 520 nm. For experiments in which binding was determined by ssDNA-induced quenching of the intrinsic tryptophan fluorescence of POT1, a fixed concentration of protein was titrated with unlabeled O1. Excitation was at 290 nm and emission at 340 nm was recorded. Kinetic experiments (described below) showed that binding of ssDNA to POT1 is fast (relaxation time t » 80 ms); thus, complex formation between POT1 and ssDNA could be followed by conventional titrations of serial additions of ligand to receptor without complications due to slow processes. Determination of accurate equilibrium dissociation constants from POT1 tryptophan emission experiments required application of an inner filter correction (2) due to the (low) absorbance of the added DNA at the excitation and emission wavelengths. K_d_ values were estimated for both types of titration by fitting the titration curve to a 1-site ligand depletion model (3) (Eq. 1) by the non-linear least squares procedure in OriginPro 2016 (OriginLab, Inc., Northhampton, MA).

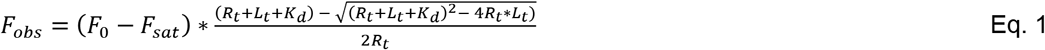

*F_obs_* is the observed fluorescence intensity, *F_0_* is the fluorescence intensity before ligand addition, *F_sat_* is the fluorescence intensity at saturation, *R_t_* = total receptor concentration, *L_t_* is the total ligand concentration at each addition, and *K_d_* is the dissociation constant. *F_0_, F_sat_*, and *K_d_* were optimized by non-linear least squares.

##### FRET-labelled G4s

Binding of POT1 to G4 DNA was slow, requiring >2 hr at room temperature to achieve equilibrium. Thus to determine POT1-G4 K_d_ values, the slow rate of equilibration required that each point in a titration be determined independently using an individual POT1:G4 mixture rather than by serial additions of POT1 to the same G4 sample. We therefore mixed FRET-labelled G4 with an appropriate concentration of POT1 followed by incubating the samples overnight in the dark at room temperature. Binding isotherms were constructed from these solutions by measuring the increase in 6FAM fluorescence brought about by G4 unfolding and POT1 binding as described above for FRET-O1.

##### Data analysis

POT1 binding isotherms were fit to the binding models shown in Table S3 using user defined functions implemented in GraphPad Prism version 6.07 (GraphPad Software, Inc., San Diego, CA). Monte Carlo simulations used a strategy previously used in our laboratory (4), but were implemented using a tool available in GraphPad Prism.

#### Kinetic Experiments

##### Stopped-flow kinetics

The kinetics of POT1-FRET-O1 interaction was assessed by stopped-flow mixing using an OLIS RSM-SF instrument (OLIS Instruments, Bogart, GA) in the fluorescence mode. The drive syringes and observation cuvette were maintained at 25 °C with a circulating water bath. The 6FAM label of the O1 was excited at 490 nm and the increase in emission subsequent to mixing was measured at right angles to excitation through a 520 nm interference filter. The kinetic data were fit to a single exponential using the non-linear least squares routine in the program OriginPro 2016 to obtain best-fit signal amplitudes and relaxation times.

##### G4 unfolding kinetics

As previously described (5), unfolding of FRET-G4s is slow and accompanied by an increase in 6FAM emission intensity that accurately tracks the progress of the unfolding reaction. Briefly, a 5-fold excess of either complementary DNA or POT1 was added to pre-folded G4 in POT1 buffer and manually mixed. The kinetics of G4 unfolding was followed at 25 °C using the SpectraMax-3 fluorometer with excitation at 490 nm and emission at 520 nm. The kinetic profile of unfolding in the presence of complementary DNA or POT1 required three exponentials with relaxation times of ~10^2^, ~10^3^ and ~10^4^ s for accurate fitting. The optimized amplitudes and relaxation times were determined by non-linear least squares regression with OriginPro 2016 as previously described (5).

#### Isothermal Titration Calorimetry (ITC)

The ITC titrations were obtained using a Microcal VP-ITC microcalorimeter (Microcal, Northhampton, MA). The O1 oligonucleotide was dissolved in dH_2_O then diluted into 20mM Potassium Phosphate, 180mM KCl pH 7.2, to a final concentration at 56uM. POT1 was dialyzed overnight in POT1 buffer then diluted to 2.6uM. ITC titrations were done at 25°C with 5μL injections, duration 10 seconds, spacing 240 seconds and filter at 5. The reference power was 15μcal/seconds, stirring speed 300 rpm and 60 second initial delay. Data processing and analysis were done using Affinimeter software v. 1 (AFFINImeter, Santiago de Compostela. Spain; https://www.affinimeter.com/site/).

#### Analytical Ultracentrifugation

Sedimentation velocity experiments were carried out in a Beckman Coulter ProteomeLab XL-A analytical ultracentrifuge (Beckman Coulter Inc., Brea, CA) at 20°C and 50,000 rpm in standard 2 sector cells. Buffer density was determined on a Mettler/Paar Calculating Density Meter DMA 55A at 20.0 °C and viscosity was measured using an Anton Parr AMVn Automated Microviscometer at 20°C. Data were analyzed with the program Sedfit (free software: www.analyticalultracentrifugation.com) using the continuous c(s) distribution model. The partial specific volume of POT1 was calculated from its amino acid composition (0.7453 ml/g) using the Protparam tool in ExPASy (free software: web.expasy.org) and a value of 0.55 ml/g was used for the DNA oligonucleotides (14). The stoichiometry of DNA:POT1 complexes was determined using the two wavelength method described by Brautigam et al. (6). POT1-oligonucleotide mixtures were scanned at both 260 and 280 nm during centrifugation and the absorbance at both wavelengths in the largest species was determined using integration of the peak in the c(s) vs s mode of sedfit. The concentration of each species in the peak could be determined and the stoichiometry obtained using the extinction coefficients of POT1 and oligonucleotide at both wavelengths. The partial specific volumes for protein:DNA complexes were calculated from the weight averages of each component in the complex (7) and used to determine the molecular weight from c(s) vs s analysis. Frictional ratios, f/fo, were determined using Ultrascan3 (free software: uslims3.uthscsa.edu), since f/fo reported by Sedfit analysis is the weight average value for all species present. Calculations were performed on the UltraScan LIMS cluster at the Bioinformatics Core Facility at the University of Texas Health Science Center at San Antonio and multiple High Performance Computing clusters supported by NSF XSEDE Grant #MCB070038 (to Borries Demeler).

#### Molecular Dynamics Simulations

The structural models used in molecular dynamics simulations were hPOT1 (PDB ID: 1XJV) and 23-nt hybrid-1 telomeric G-quadruplex (PDB ID: 2GKU). The unmodified 2GKU structure was used. For POT1 (1XJV), residues PRO146, SER147, and TRP148 were added between OB1 and OB2 in Maestro 11.8 (Schrödinger). This complete POT1 structure was subjected to a total of 100 ns molecular dynamics production trajectories (as described below) with and without the bound 10mer oligonucleotide O1 (Table S1), allowing them to relax and explore conformational space. The POT1:O1 complex was subsequently used to manually construct the 2GKU-POT12 complex. This was accomplished by loading two copies of the relaxed POT1-O1 complex into Maestro and manually connecting them using a guanine dinucleotide d(GG). This was followed by the removal of the 5’ dT from the POT1 on the 5’-end, mutation of the 5’-most dA to dT, and addition of a dA to the 3’ end to match the 2GKU sequence. This complex was subsequently saved as a PDB for further simulation. Last, the 2GKU-duplex was constructed using Maestro in a B-form DNA conformation.

Molecular dynamics simulations were carried out on the following models: POT1 alone, 2GKU-POT12, and 2GKU-duplex. The PDB structures were imported into the xleap module of AMBER 2018, neutralized with K+ ions, and solvated in a rectangular box of TIP3P water molecules with a 15 Å buffer distance. All simulations were equilibrated using sander with the following steps: (1) minimization of water and ions with restraints of 10.0 kcal/mol/Å on all nucleic acid and amino acid residues (2000 cycles of minimization, 500 steepest decent before switching to conjugate gradient) and 10.0 Å cutoff, (2) heating from 0K to 100K over 20 ps with 50 kcal/mol/Å restraints on all nucleic acid and amino acid residues, (3) minimization of entire system without restraints (2500 cycles, 1000 steepest decent before switching to conjugate gradient) with 10 Å cutoff, (4) heating from 100K to 300K over 20 ps with restraints of 10.0 kcal/mol/Å on all nucleic acid and amino acid residues, and (5) equilibration at 1 atm for 100 ps with restraints of 10.0 kcal/mol/Å on nucleic acids. The output from equilibration was then used as the input file for 100 ns of unrestrained MD simulations using pmemd with GPU acceleration in the isothermal isobaric ensemble (P = 1 atm, T = 300 K). Periodic boundary conditions and PME were used. 2.0 fs time steps were used with bonds involving hydrogen frozen using SHAKE (ntc = 2). Trajectories were analyzed using the cpptraj module in the AmberTools 18 package.

Hydrodynamic properties were calculated as average and standard deviation using equally spaced snapshots across the entire trajectory, or the single PDB file in the case of the G4 2GKU. This was accomplished using HYDROPRO10. For the G4 2GKU and 2GKU-duplex structures an atomic level calculation was performed (INMODE=1, AER = 2.53) with vbar = 0.55. For POT1 and 2GKU-POT12, a residue level calculation was performed (INDMODE=2, AER=4.8) with vbar = 0.73 and 0.712, respectively. The vbar for the protein-DNA complex was modified by weighting to account for the vbar contribution of DNA using the equation vbar_weighted_ = ((0.73*a) + (0.55*b)), where *a* and *b* are the weight fractions of protein and nucleic acid in the complex, respectively. All HYDROPRO calculations used temperature of 20.0°C, viscosity = 0.0101 poise, and density = 1.0092 g/cm^3^. Molecular weights were the same as given in Table 2, rounded to the thousandth place

**Table S1.**
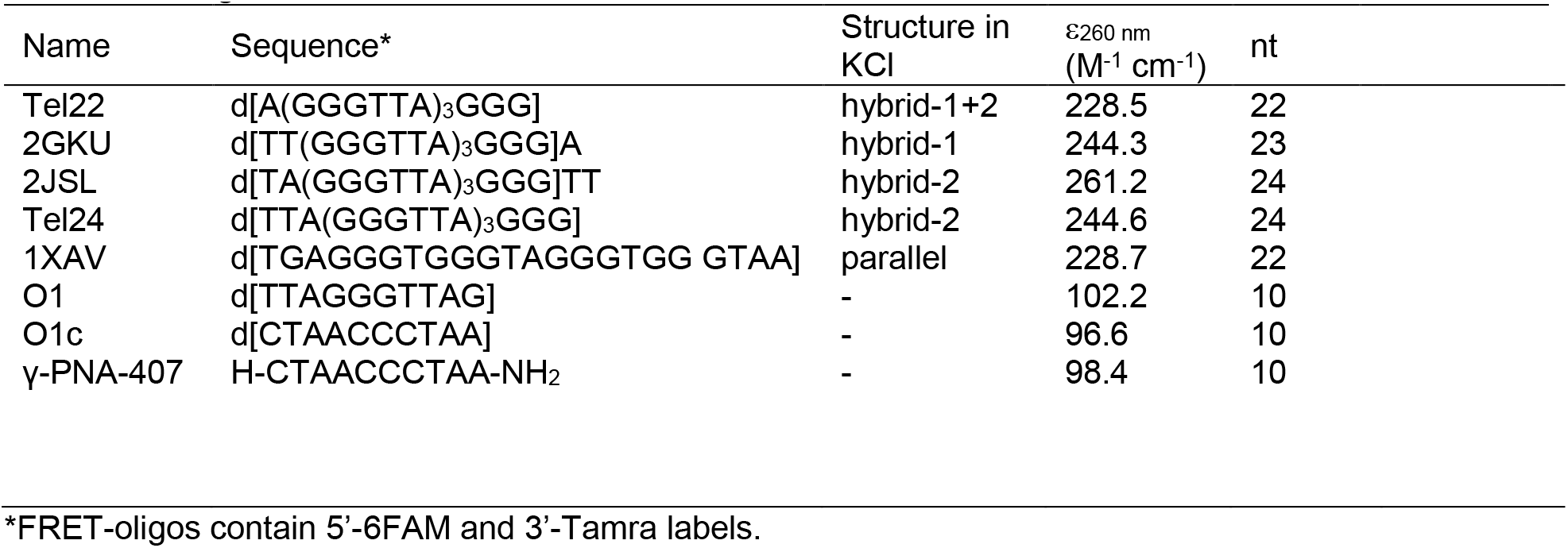
Oligonucleotides used.

**Table S2.** Summary of binding constants for O1-POT1 interaction.

| Method | $K_d$ nM | $K_a/10^7 \text{ M}^{-1}$ | $\Delta G_{25^\circ\text{C}}$ kcal/mol |
| --- | --- | --- | --- |
| FRET | $26.4 \pm 17.0$ | 3.8 | -10.3 |
| POT1 Trp | $29.6 \pm 36.0$ | 3.4 | -10.2 |
| MST* | $58.5 \pm 12.3$ | 1.7 | - 9.8 |
| 2-AP | $17.2 \pm 8.7$ | 5.8 | -10.6 |
| ITC | $59.4 \pm 3.0$ | 1.68 | -9.8 |
| Average | $38.2 \pm 19.5$ | $3.3 \pm 1.7$ | $-10.1 \pm 0.3$ |

All experiments done in 20 mM K Phosphate, 180 mM KCl, pH 7.2, 25 C.
\*MST titration done in 20 mM K Phosphate, 20 mM KCl, pH 7.2, 25 C.

**Table S3.** Comparison of fitting models for the POT1-G4 binding isotherm.

| Model | $K_d$ μM | $Y_{max}$ | n | $K_c$ | ASSR | Sy.x | P value |
| --- | --- | --- | --- | --- | --- | --- | --- |
| **One Site** | 0.95 ± 0.2 | 1.84 ± 0.2 | - | - | 0.1725 | 0.0619 | |
| **Hill** | 0.32 ± 0.02 | 1.01 ± 0.04 | 1.88 ± 0.13 | - | 0.0614 | 0.0374 | <0.0001 |
| **CS** | 0.02 ± 0.03 | 0.98 ± 0.04 | 2.13 ± 0.037 | 0.003 ± 0.006 | 0.0604 | 0.0375 | <0.0001 |
| **CS**<br>**Fixed $K_d$** | (0.038) | 0.96 ± 0.03 | 2.32 ± 0.14 | 0.006 ± 0.002 | 0.0608 | 0.0372 | - |
| **CS**<br>**Fixed n** | 0.012 ± 0.01 | 0.99 ± 0.02 | (2.0) | 0.0012 ± 0.002 | 0.0606 | 0.0371 | - |
| **CS**<br>**Fixed $K_c$** | 0.011 ± 0.009 | 0.99 ± 0.04 | 2.02 ± 0.16 | (0.00114) | 0.0606 | 0.0371 | - |

**Figure S1.**
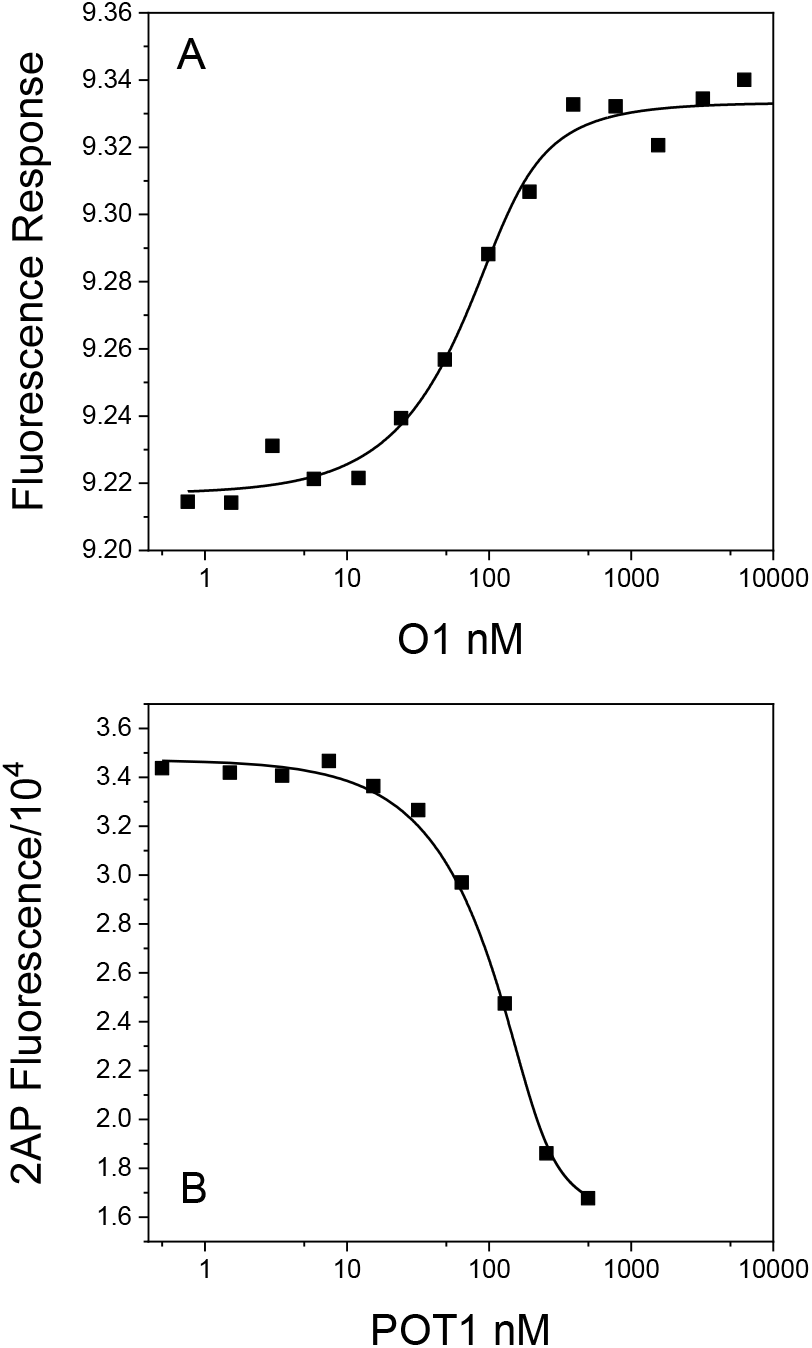
Binding isotherms for the interaction of POT1 and O1. (A) Data obtained by microscale thermophoresis (MST) with 25 nM NT-647 labeled POT1 and increasing amounts of added O1. (B) Data obtained by fluorescence with 200 nM 2-aminopurine labeled O1 and increasing amounts of added POT1. The solid lines are the best fits to a 1:1 binding model with the K_d_ values shown in Table S1.

**Figure S2.**
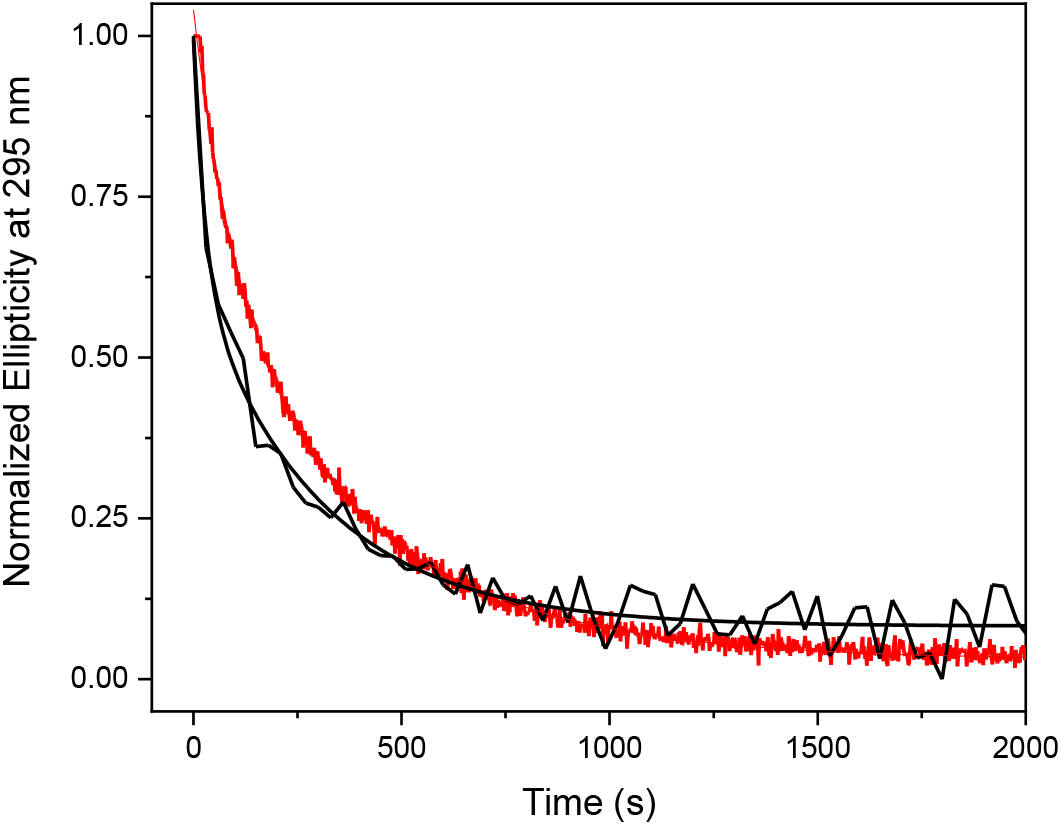
Kinetics of Tel22 unfolding monitored by circular dichroism at 295 nm. The black curve shows unfolding induced by addition of 3.7 μM POT1 to 1.2 μM Tel22. The red curve shows unfolding by addition of 5 μM Tel22 complement to 4.8 μM Tel22. Conditions: 20 mM sodium phosphate, 180 mM NaCl, pH 7.2, 25 °C.

**Figure S3.**
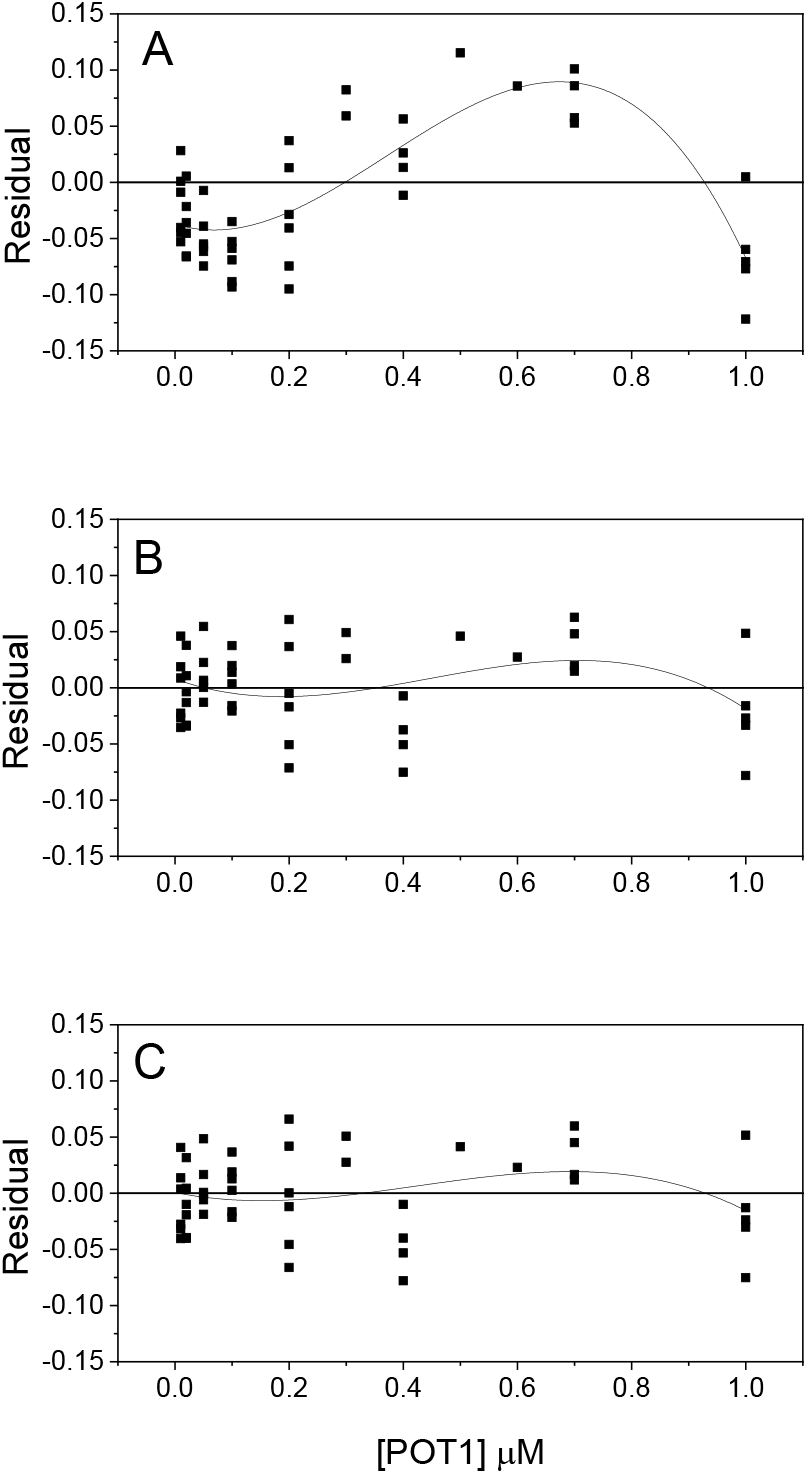
Residual plots for fits to the POT1-G4 DNA binding isotherm. (A) Simple, noncooperative binding (B) The Hill equation (C) The conformational selection model

**Figure S4.**
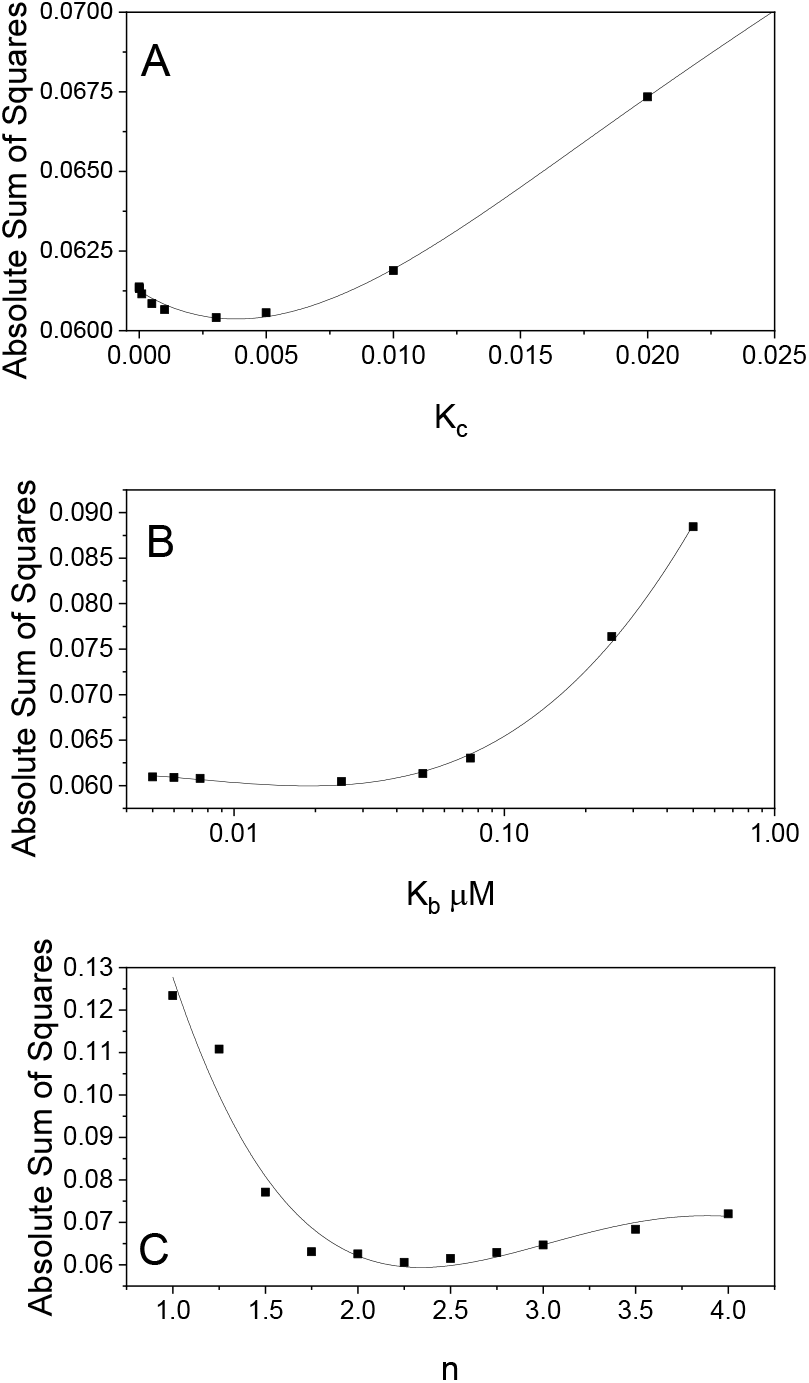
Error spaces for fitting parameters in the conformational selection model. (A) Error as a function of K_c_, the G4 unfolding equilibrium constant. (B) Error as a function of the POT1 binding constant K_b_. (C) Error as a function of n, the binding stoichiometry.

**Figure S5.**
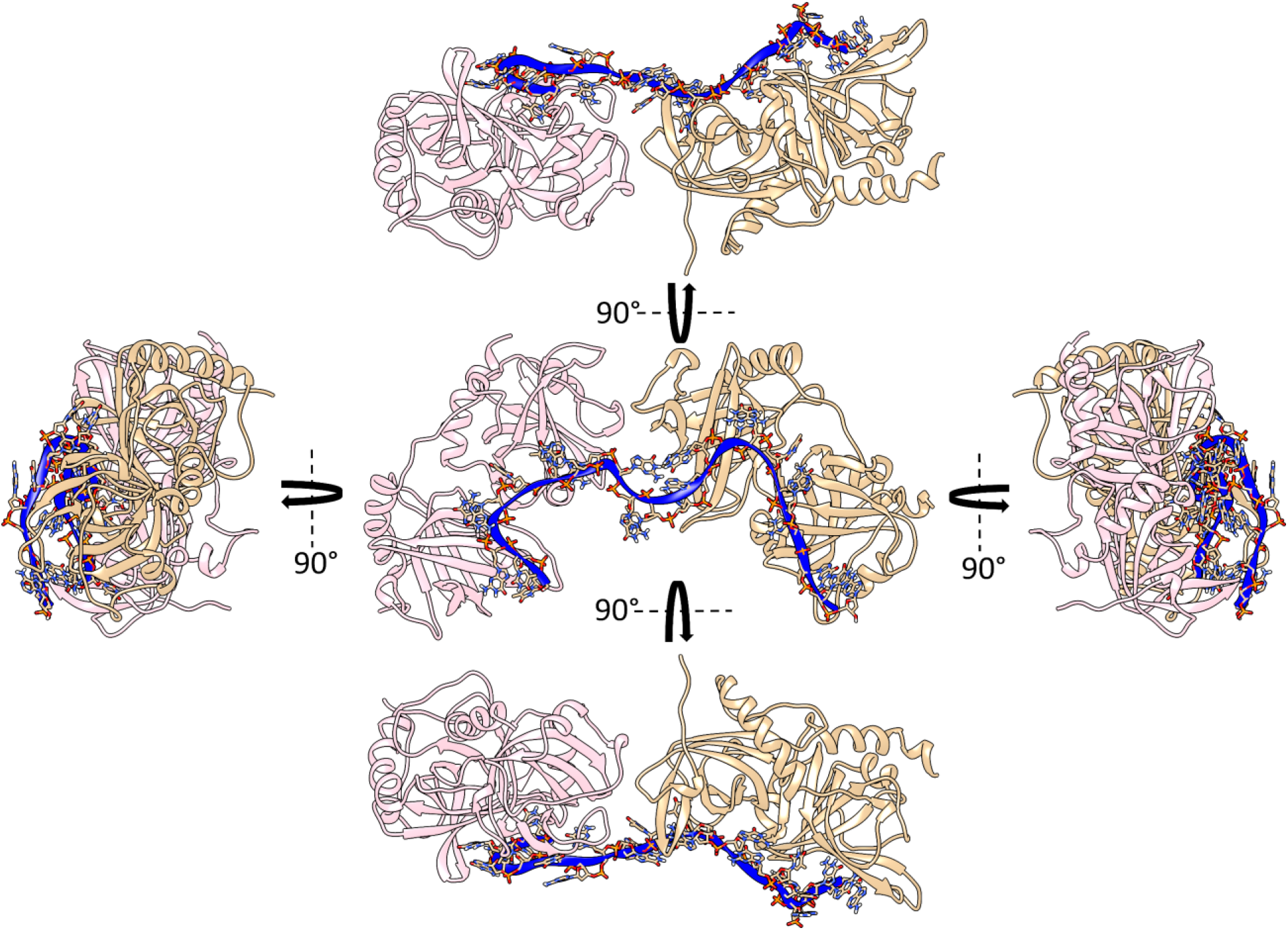
Orthogonal views of the POT1-telomeric DNA complex. The DNA strand is colored in blue. Two POT1 molecules, colored separately, are bound to the DNA.

**Figure S6.**
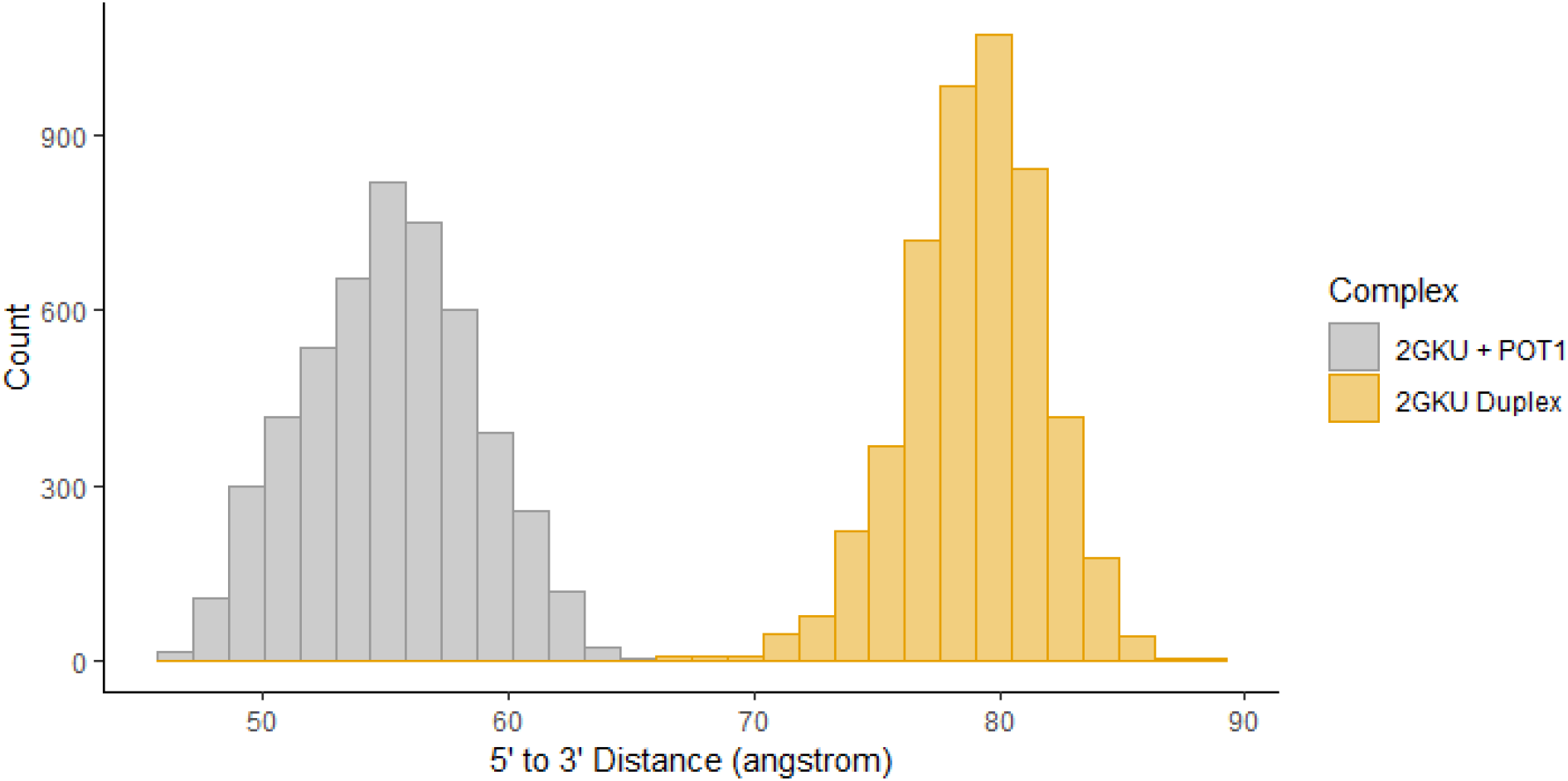
Distributions of the end-to-end distances of 2GKU DNA in duplex form (gold) or as a single-strand in a 2:1 POT1:DNA complex (gray). The 5’-OH and 3’-OH oxygen atoms were used for end-to-end distances for the DNA in both complexes were measured over the entire 10 ns trajectory of the simulation, yielding estimates of 55.2 ±3.5 Å and 78.9±2.9 Å for the POT1 complex and duplex form, respectively.

